# Telomeric Double Strand Breaks Facilitate Formation of 5’ C-Rich Overhangs in G1 Human Cells

**DOI:** 10.1101/720565

**Authors:** Christopher B Nelson, Taghreed M Al Turki, Lynn Taylor, David G Maranon, Keiko Muraki, John P. Murnane, Susan M Bailey

## Abstract

Telomeres are repetitive nucleoprotein complexes that protect chromosomal termini and prevent them from activating an inappropriate DNA damage response (DDR). Here, we characterized the human cellular response to targeted telomeric DSBs in telomerase positive and telomerase-independent alternative lengthening of telomeres (ALT) cells, specifically in G1. Telomeric DSBs in G1 human cells elicited early signatures of a DDR, however, localization of 53BP1, an important regulator of resection at broken ends, was not observed at telomeric break sites. Consistent with this finding and previously reported repression of classical nonhomologous end-joining (c-NHEJ) at telomeres, evidence for c-NHEJ was also lacking. Likewise, no evidence of homologous recombination (HR)-dependent repair of telomeric DSBs in G1 was observed. Rather, and supportive of rapid truncation events, telomeric DSBs in G1 human cells facilitated formation of extensively resected tracks of 5’ C-rich telomeric single-stranded (ss)DNA, a previously proposed marker of the recombination dependent ALT pathway. Indeed, induction of telomeric DSBs in human ALT cells also resulted in significant increases in 5’ C-rich (ss)telomeric DNA in G1, which rather than RPA, were bound by the complementary telomeric RNA, TERRA. These results suggest that targeting TERRA-mediated protection at damaged telomeres may represent a promising therapeutic strategy, particularly against ALT-positive cancers.

## INTRODUCTION

Telomeres, specialized nucleoprotein complexes that “cap” the ends of linear chromosomes, are composed of highly conserved, G-rich tandem repeats [(5’-TTAGGG-3’)n in vertebrates] (1). Due to their repetitive nature and abundance of heterochromatic marks, telomeres were long regarded as silenced, non-transcribed features of the genome. Thus, the relatively recent discovery of Telomere Repeat-containing RNA (TERRA) opened many new avenues of investigation (2). TERRA is a long, noncoding RNA (lncRNA) shown to serve a structural role at telomeres, as well as function in regulation of telomere length and telomerase activity, thus TERRA participates in maintenance of telomeres and genome stability (3–7).

Telomeres end with a 3’ single-stranded (ss)G-rich overhang (8), which serves as the substrate for telomerase-mediated synthesis of telomeric DNA (9). Telomerase is the reverse transcriptase capable of maintaining telomere length via RNA template-dependent addition of telomeric repeats onto the ends of newly replicated chromosomes. Telomerase activity is prominent in highly proliferative populations like germ-line, stem, and the vast majority of cancer cells, thereby endowing them with extended or unlimited replicative potential (10, 11). The remaining ~10% of human cancers maintain telomere length via a recombination-dependent, alternative lengthening of telomeres (ALT) mechanism (12) that display a number of defining features, including heterogeneous telomere lengths, increased frequencies of telomere sister chromatid exchange (T-SCE), ALT-associated PML bodies (APBs), and extrachromosomal telomeric repeats (ECTR), which include C-rich (ss)circles (C-circles) (13–16). The ALT phenotype is relatively common in several subtypes of human sarcomas and astrocytomas, and has been observed in ~4% of all tumor types, including carcinomas and pediatric glioblastoma multiformes (17).

The telomeric 3’ (ss)G-rich overhang is also required for the formation of protective terminal structural features termed T-loops (18). Telomeres are bound shelterin, a protein complex that includes the telomere-repeat binding factors TRF1 and TRF2, which contribute to regulation of telomerase activity, T-loop formation, and protection of chromosome ends (19). Functional telomeres are essential for maintaining genome stability, as they protect natural chromosomal termini from degradation and prevent them from being recognized as double strand breaks (DSBs) and triggering of an inappropriate DNA damage response (DDR) (20–23). Inhibition of conventional repair activities at telomeres has been demonstrated (22, 24–26), raising the question of how – and even whether – DSBs occurring within telomeric repeats themselves are repaired. Various strategies employing targeted enzymatic cleavage of telomeric repeats have recently enabled studies to directly address this intriguing issue (27–30).

Enzymatically induced telomeric DSBs in murine cells have been shown to activate a DDR and recruitment of p53 binding protein 1 (53BP1) in a subpopulation of cycling cells, specifically those undergoing DNA replication (30). Moreover, homologous recombination (HR) and alternative non-homologous end-joining (alt-NHEJ), but not classical NHEJ (c-NHEJ), occurred following induction of telomeric DSBs in cycling cell populations (29, 30). These results suggest that while repair of telomeric DSBs is possible, it may be limited to cells undergoing replication. Such a notion is also supported by studies utilizing global DNA damaging agents – ionizing radiation (IR) and hydrogen peroxide – which although would only rarely be expected to directly produce telomere-specific DSBs, have shown that telomeric damage responses persist in G1 cells that undergo senescence (31, 32).

Whether or not repair of telomeric DSBs requires cell cycle progression (replication) has physiological relevance, as many human adult tissues are largely post-mitotic and unrepaired DSBs can trigger senescence, thereby contributing to degenerative pathologies (31, 32). Here, we investigated human cellular responses to targeted telomeric DSBs specifically in G1 utilizing the previously characterized telomere-specific endonuclease TRAS1-EN-TRF1 (EN-T) (27, 28), in telomerase positive fibroblasts and cancer cells, and telomerase-independent ALT cells. Signatures of an early DDR were observed, as both gamma (γ)–H2AX and mediator of DNA damage checkpoint protein 1 (MDC1) foci co-localized with broken telomeres. However, and consistent with previous reports, 53BP1 was not recruited to telomeric DSBs in G1 (30).

Due to the scarcity of a homologous template, NHEJ is regarded as the primary DSB repair pathway during G1 in mammalian cells (33–35). Consistent with inhibition of c-NHEJ at telomeres (36, 37), we show with both short hairpin (sh)RNA depletion and chemical inhibition of the key NHEJ kinase, DNA-dependent protein kinase catalytic subunit (DNA-PKcs), that c-NHEJ is not a major contributor to repair of telomeric DSBs in G1 human cells. Likewise, no evidence of classical HR-dependent repair of telomeric DSBs in G1 was found, as neither RAD51, RAD52 (early responders that promote and stimulate strand invasion, respectively), nor repair associated DNA synthesis (BrdU incorporation) were detected. The most striking observation at telomeric DSBs in G1 were extensive tracks of predominantly 5’ C-rich (ss)telomeric DNA, which co-localized with Replication Protein A (RPA) in telomerase positive human cells. Consistent with this finding, S4/S8 phosphorylated RPA (pRPA) foci, which are associated with activation of RPA during DNA repair (38), had only modest dependence on the conventional end processing exonucleases MRE11 (3’-to-5’) and EXO1 (5’-to-3’). The 5’-to-3’ nuclease Apollo, which has been implicated in post-replicative processing specifically of leading-strand telomeres (39), also did not influence resection at telomeric DSBs in G1.

These results support the view that telomeric DSBs in G1 human cells represent rapid truncation events, in that they facilitate formation of 5’ C-rich (ss)overhangs, previously proposed markers of the recombination and replication-dependent ALT pathway of telomere maintenance (40). Indeed, induction of telomeric DSBs in human U2OS (ALT) cells also resulted in significant increases in 5’ C-rich (ss)telomeric DNA in G1, which was bound by the complementary telomeric RNA, TERRA. We propose that enrichment of 5’ C-rich (ss)telomeric DNA in G1 results from telomere DSB-mediated deletion of protective T-loops and extensive resection in the absence of 53BP1. Furthermore, these exposed and vulnerable structures are protected by interactions involving RPA and transient telomeric RNA: DNA hybrids (41), which are dependent on telomerase/ALT status, and presumably allow them to persist into S/G2 for efficient replication/HR-dependent elongation, while also circumventing conventional repair pathways (42). Thus, targeting TERRA-mediated protection at damaged telomeres may represent a promising therapeutic strategy, particularly against ALT-positive cancers.

## RESULTS

### Characterization of targeted telomeric DSBs in human G1 cells

To better understand human cellular responses to telomeric DSBs throughout the cell cycle, we performed transient transfection experiments using a previously characterized plasmid encoding a flag-tagged telomere repeat-specific endonuclease fused to the human TRF1 gene (TRAS1-EN-TRF1: hereafter referred to as EN-T) that produces blunt ended DSBs within telomeres (27, 28). Several human cell lines with different telomerase status were selected. U2OS (ALT, telomerase independent) cells served as a positive control for EN-T activity as cycling ALT cells undergo recombinational repair at telomeric DSBs. BJ1 hTERT immortalized fibroblasts (telomerase positive) represent an apparently normal, non-tumorigenic human cell line. Lastly, EJ-30, a bladder carcinoma cell line, was employed (highly telomerase positive), as they have been used extensively to study sub-telomeric DSB repair (36, 37, 43).

Following transient transfection with EN-T or TRF1-only, co-localization with telomeres was observed in all cell lines (**Supp Fig 1A**). Evidence of DSB signaling following EN-T expression in EJ-30 (non-ALT, cancer) cells was also evaluated, which included phosphorylation of ATM (S1981) and CHK2 (Thr68) (**Supp Fig 1B**). Consistent with previous reports of TRF1 overexpression inducing DSBs [although via a different mechanism involving telomere association, anaphase bridges and breakage (44–46)], TRF1-only also induced DSB signaling activity, noted by the increase in intensity of phospho-ATM and phospho-CHK2 bands in transfected samples relative to no treatment controls. Supportive of telomere-specific cutting and rapid truncation events with EN-T, a modest decrease in Telomere Restriction Fragment (TRF) size was also observed following EN-T expression, as compared to TRF1-only and untransfected controls (**Supp Fig 1C, D**). EN-T expression reduced the mean TRF by ~ 5-10%, consistent with expectation considering the relatively low transfection efficiency in EJ-30 cells (~20-30%), and that not all telomeres were broken (~8-12/cell visualized).

To further validate the EN-T system, we sought to reproduce the finding that induced telomeric DSBs stimulate a damage response and repair via some combination of HR and break-induced replication (BIR) in cycling ALT cells (47, 48). Human U2OS (ALT) cells exhibited activation of telomere damage responses upon transfection with EN-T as expected, and as evidenced by increased γ-H2AX foci compared to untransfected controls. Importantly, a significant portion of these well-accepted DSB damage markers occurred at broken telomeres, as γ-H2AX foci colocalized (overlapped) with EN-T foci (**Supp Fig 2A**). Additionally, following EN-T transfection, cycling U20S cells harbored elevated levels of RAD51 and RAD52-YFP foci, mediators of HR and BIR respectively, which frequently colocalized with EN-T foci, confirming repair of telomeric DSBs by HR and BIR in cycling human ALT cells (**Supp Fig 2B, C**).

As various lines of evidence have suggested that DSB repair may be non-conventional or even non-existent within or near telomeres in G1 (29–32), we sought to investigate DNA damage responses and repair of broken human telomeres in G1 directly. In order to study telomeric DSBs specifically in G1 non-ALT EJ-30 (telomerase positive; cancer) cells, we employed a DAPI intensity-based approach as a means of distinguishing cell cycle phases in interphase nuclei, which retained the ability to make accurate measurements of fluorescent foci. Cells in G1 form a clear peak in the lower intensity portion of a DAPI intensity histogram using even a relatively low number (~300) of cells (**Supp Fig 3A**). The specificity of the G1 DAPI intensity peak was validated via exclusion of Cyclin A, which stains S and G2 cells. A similar DAPI intensity histogram was generated to distinguish G1 from S/G2 in all imaging experiments involving EJ-30 cells. Reliable discrimination between S and G2 could not be achieved with this approach, therefore these populations were pooled throughout analyses.

Although transfection efficiencies in BJ1 hTERT cells were quite low (0.5-2%), telomeric DSB induction was evaluated in EN-T and TRF1-only transfected cells. Since only a very small percentage of transfected cells stained positive for Cyclin A (EN-T: 0%, TRF1: 3.2%**, Supp Fig 3B**) or incorporation of bromodeoxyuridine (BrdU; **Supp Fig 3C**), the vast majority of transfected BJ1 hTERT cells were in G1 phase 48 hr post transfection. Transfection efficiencies were much higher (~25-30%) in U2OS (ALT) cells (EN-T, TRF1-only, empty vector). A stably transfected Fluorescent Ubiquitination-based Cell Cycle Indicator (FUCCI) (49) U2OS cell line was also generated to definitively identify cell cycle phase, and for these EN-T experiments, scoring was restricted to only transfected cells (100%).

### Non-canonical damage response at telomeric DSBs in G1 human cells lacks recruitment of 53BP1

I-SCE1 induced DSBs in sub-telomeric regions have previously been shown to be deficient in DSB repair (36, 37). To investigate damage responses at DSBs within individual telomeres, we evaluated co-localization of γ-H2AX and 53BP1 foci at broken telomeres. Telomere-specific DSBs co-localized with γ-H2AX in BJ1 hTERT G1 cells (p = 0.012; **Fig 1A**), and in all phases of the cell cycle in EJ-30 cells (p = 0.0009 in G1 cells; p = 0.022 in S/G2 cells; **Fig. 1C**). Telomeric DSBs also co-localized with 53BP1 in EJ-30 S/G2 cells (p = 0.012); however, 53BP1 was not observed at broken telomeres in either BJ1 hTERT or EJ-30 G1 cells (**Fig 1B, D**). To determine whether other components of the early DNA damage response were activated by telomeric DSBs in G1, we also evaluated MDC1, an early mediator of the response to genomic DSBs that acts downstream of γ-H2AX, but upstream of 53BP1 (50). MDC1 foci were induced to a similar degree as γ-H2AX in response to telomeric DSBs in BJ1 hTERT G1 cells (p = 0.0007, **Fig 2A**).

**Figure 1:**
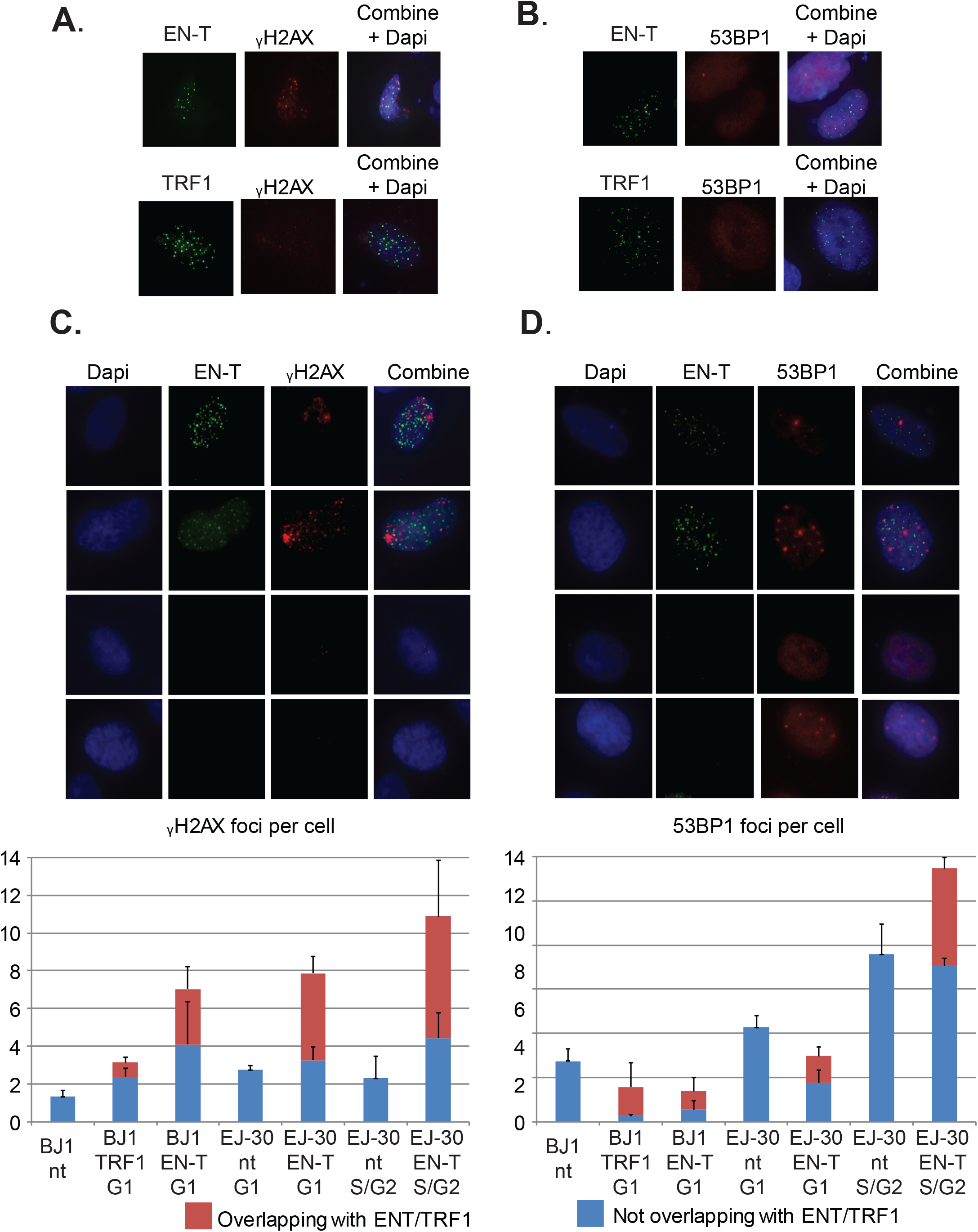
Telomeric DSBs co-localize with γ-H2AX, but do not attract 53BP1 to break sites in G1. **A.** γ-H2AX foci co-localized with broken telomeres following transfection with EN-T in BJ1 hTERT G1 cells (p=0.012). **B.** However, 53BP1 foci did not co-localize with EN-T induced telomeric DSBs in BJ1 hTERT G1 cells. **C, D**. EJ-30 cells displayed similar G1 DNA damage response as BJ1 hTERT cells. While telomere DSB induction by EN-T stimulated γ-H2AX foci at telomeres in both G1 and S/G2 cells (p = 0.0009 and 0.022 respectively), 53BP1 foci were only detected at telomeres in S/G2 cells (p = 0.012). Data represent three independent experiments (n= 30 BJ1 hTERT; n= 300 EJ-30 cells/experiment). Error bars are SD, p values determined using ANOVA with Tukey’s post hoc test for significance; p <0.05 is significant.

**Figure 2:**
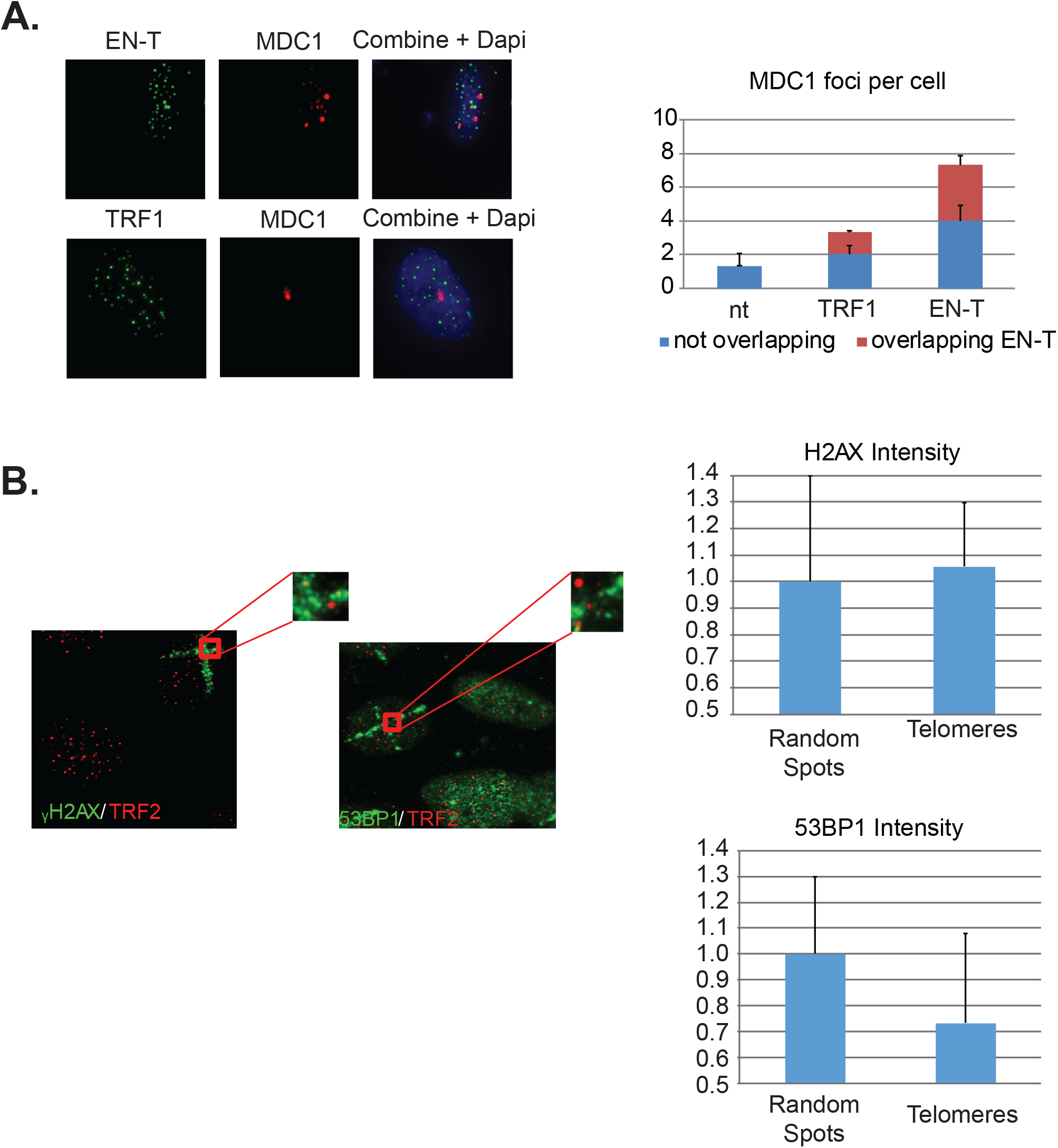
DNA damage response at telomeric DSBs in G1. **A.** Transfection with EN-T also resulted in an increased number of MDC1 foci in BJ1 hTERT G1 cells (compared to non-transfected and TRF1 control cells), which often overlapped with ENT-flag (p = 0.0007). Data represent three independent experiments (n=30). Error bars are SD, p values determined using ANOVA with Tukey’s post hoc test for significance; p <0.05 is significant. **B**. The intensity of γ-H2AX within microirradiation-induced DNA damage stripes was similar at telomeres and random spots within non-transfected BJ1 hTERT cells, while the intensity of 53BP1 within damage stripes was decreased at telomeres relative to random spots (p=0.099).

To evaluate whether other DNA damaging methodologies might also initiate a response lacking 53BP1 at telomeres, we compared the intensity of γ-H2AX and 53BP1 foci co-localized at telomeres versus at random genomic sites that occurred within spatially defined stripes of damage generated by laser microirradiation. Consistent with our results using EN-T, 30 minutes after exposure of BJ1 HTERT cells, the intensity of γ-H2AX was found to be similar at telomeres and random sites within the microirradiation stripe, while the intensity of 53BP1 was reduced at broken telomeres compared to random sites (p = 0.099; **Fig 2B**).

Lastly, we reasoned that normal telomere protection activity might prevent recruitment of 53BP1 to broken telomeres. Therefore, a variety of strategies were employed to compromise telomeric end-protection, including relaxation of chromatin utilizing the histone deacetylase inhibitor Trichostatin A (or exposure to a hypotonic solution; not shown), partial depletion of the shelterin component TRF2 via small interfering (si)RNA knockdown (above the level that induces a damage response), as well as small hairpin (sh)RNA knockdown of shelterin-associated DNA-PKcs (**Supp Fig 4A-D**). However, none of these conditions resulted in recruitment of 53BP1 to telomeric DSBs in G1 human cells. Together, these results further support the finding that although telomeric DSBs in G1 activate an early DDR (γ-H2AX, MDC1 recruitment), they do not attract 53BP1 to telomeric break sites.

### c-NHEJ does not significantly contribute to repair of telomeric DSBs in human cells

The absence of 53BP1 recruitment to telomeric DSBs in G1, particularly with shRNA knockdown of DNA-PKcs, suggested that consistent with previous reports (51–53), c-NHEJ may not be occurring at broken telomeres. Autophosphorylation of DNA-PKcs at serine 2056 was slightly increased in EJ-30 cells expressing EN-T compared to cells expressing TRF1-only or no treatment controls; ionizing radiation (IR)-induced DNA-PKcs autophosphorylation was prevented by treatment with the specific kinase inhibitor NU7026 (**Fig 3A**). Considering that EN-T produced the most telomere-specific damage (**Fig 1**), we tested whether DNA-PKcs autophosphorylation influenced telomere DSB repair by comparing TRF blots in cells expressing EN-T to those expressing EN-T and treated with NU7026 (**Fig 3B**). Chemical inhibition of DNA-PKcs autophosphorylation (NU7026, 24 hr) in cycling EJ-30 cells expressing EN-T did not change the TRF size relative to the non-treated control, supporting the supposition that c-NHEJ does not significantly contribute to repair of telomeric DSBs.

**Figure 3:**
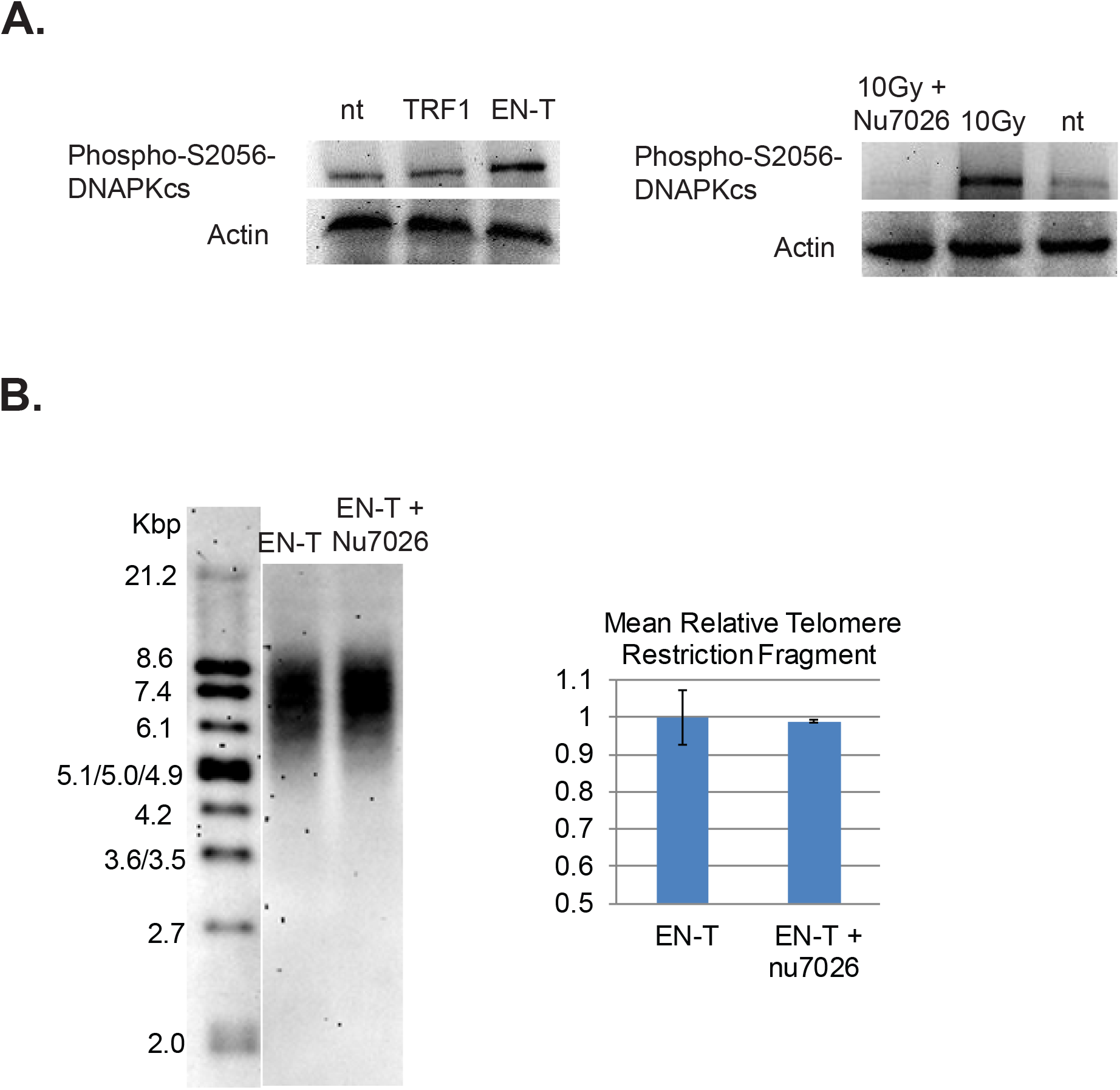
Consistent with absence of 53BP1, evidence of c-NHEJ at telomeric DSBs was lacking. **A.** Autophosphorylation of DNA-PKcs at S2056 was induced following EN-T transfection of EJ-30 cells. DNA-PKcs autophosphorylation following exposure to 10Gy ionizing radiation (gamma-rays) was prevented by the specific kinase inhibitor NU7026. **B.** To assess the role of c-NHEJ specifically at broken telomeres, EJ-30 cells transfected with EN-T were treated with NU7026, which did not significantly influence mean TRF length compared to untreated control. Error bars are SD, p values determined using ANOVA with Tukey’s post hoc test for significance; p <0.05 is significant.

### Telomere-specific DSBs in G1 are characterized by 5’ C-rich (ss)telomeric DNA

We also hypothesized that telomeric DSBs that fail to recruit 53BP1 may be particularly vulnerable to resection. To investigate the presence of ssDNA at telomeric DSBs in G1, fluorescence in situ hybridization (FISH) using a C-rich telomere probe – without denaturation of the DNA duplex [to detect 3’ G-rich (ss)telomeric DNA] – was performed in BJ1 hTERT G1 cells transfected with EN-T or TRF1-only. Indeed, telomeric ssDNA was more abundant in cells transfected with EN-T as compared with TRF1-only or no treatment controls (p = 0.0002, **Fig 4A**). To determine whether resection occurred bidirectionally, we also performed the ssFISH assay with a G-rich telomere probe [to detect 5’ C-rich (ss)telomeric DNA]. Interestingly, hybridization with the G-rich probe produced many more signals overall, and more signals in EN-T than in TRF1-only transfected cells or no treatment controls (p = 0.045, **Fig 4B**). These results revealed that the telomeric ssDNA present at telomere-specific DSBs in G1 is enriched for 5’ C-rich (ss)telomeric DNA.

**Figure 4:**
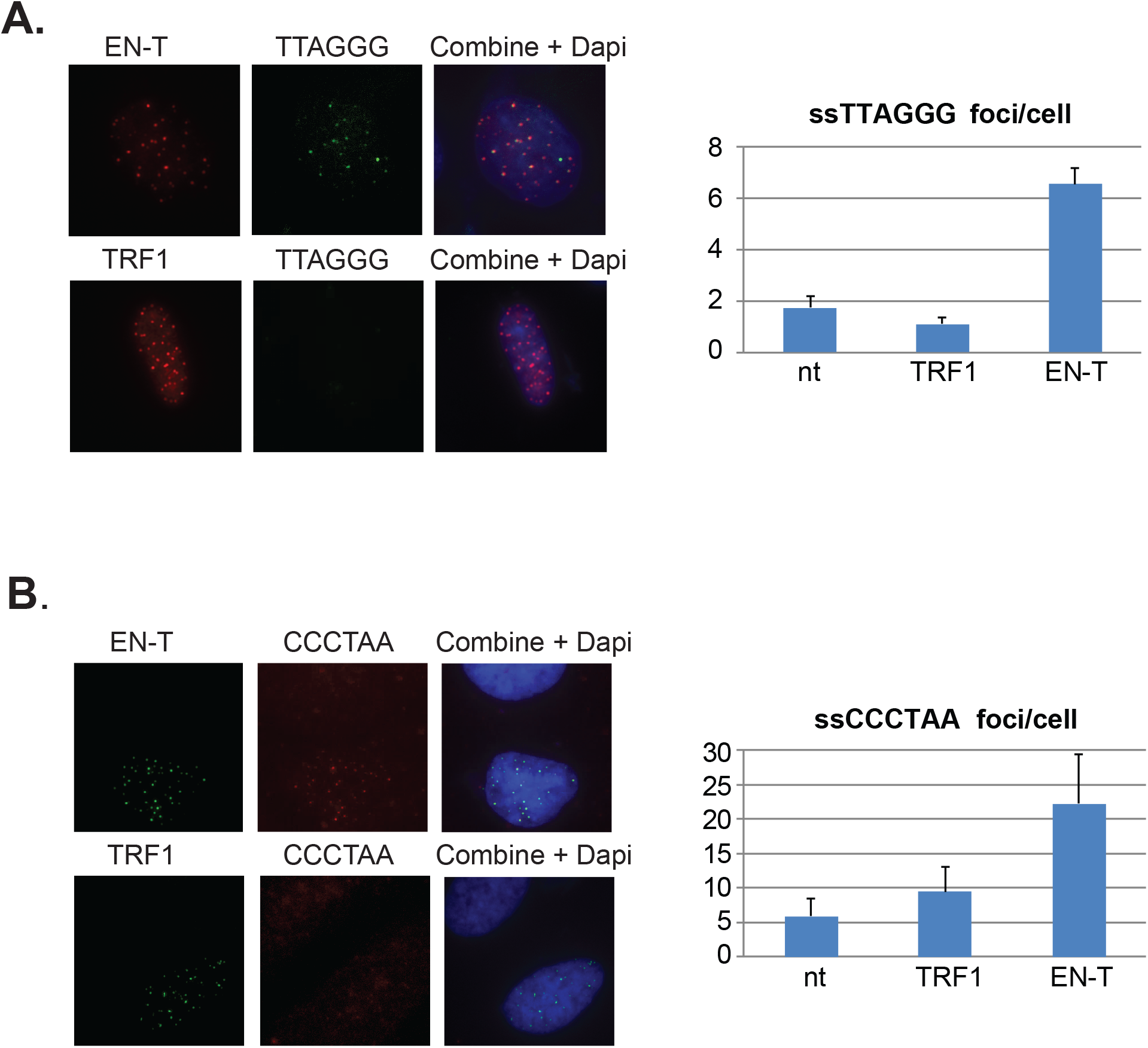
Extensive resection at telomeric DSBs in G1 facilitates formation of 5’ C-rich (ss)telomeric DNA. **A.** Transfection of BJ1hTERT cells with EN-T promoted modest production of (ss)telomeric DNA in G1 at the G-rich telomere (5’-TTAGGG-3’) (p = 0.0002). **B.** Consistent with bidirectional resection at telomeric DSBs, (ss)telomeric DNA was also associated with the C-rich telomere (5’-CCCTAA-3’) (p = 0.045), which importantly, was present at much higher frequencies. Data represent three independent experiments (n=30). Error bars are SD, p values determined using ANOVA with Tukey’s post hoc test for significance; p <0.05 is significant.

To further validate the presence of ssDNA in cells transfected with EN-T, we immunostained for RPA70 and phospho-RPA32 (S4/S8). Following induction of telomere-specific DSBs, phospho-RPA32 showed pronounced and frequent colocalization with EN-T in BJ1 hTERT G1 cells (p = 0.0000046 **Fig 5A**); RPA70 foci were not significantly increased by expression of EN-T (p = 0.27 **Fig 5B**). We observed similar increases in (ss)telomeric DNA and phospho-RPA32 in EJ-30 G1 cells expressing EN-T; as expected, this increase was also seen in S/G2 EJ-30 cells expressing EN-T (**Supp Fig 5B**).

**Figure 5:**
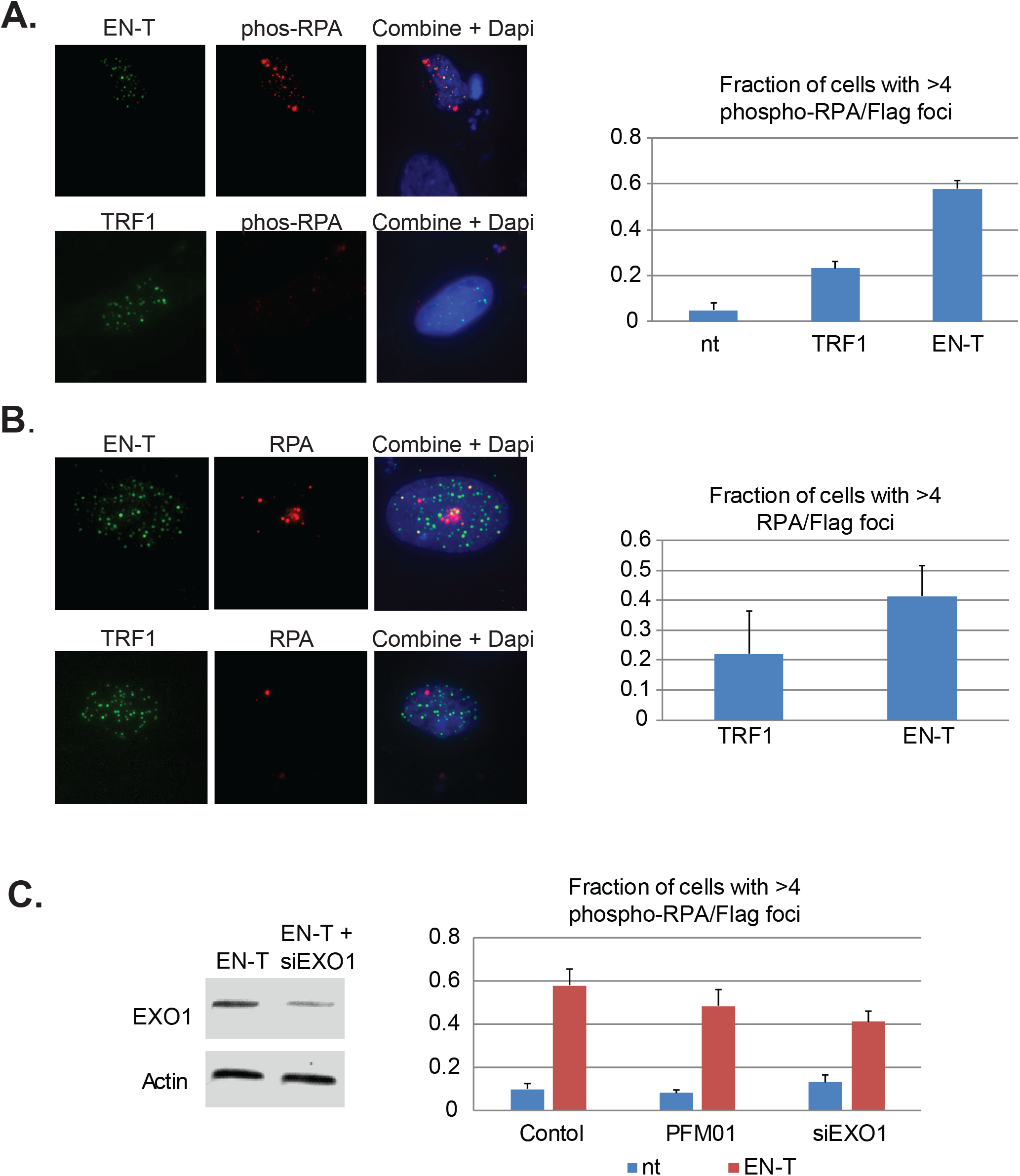
Extent of RPA-coated ssDNA at telomeric DSBs in G1 BJ1 hTERT cells is not significantly influenced by MRE11 or EXO1 nuclease activity. **A.** Transfection of BJ1 hTERT cells with EN-T induced both phospo-RPA32 and **B.** RPA70 foci in G1 that overlapped with ENT-flag (phospho RPA-32 p = 0.0000046, RPA70 p = 0.27.). **C**. Phospho-RPA32 induction following EN-T transfection was not significantly reduced by either inhibition of MRE11 endonuclease activity (PFM01) or siRNA knockdown of EXO1 (p = 0.24, 0.057, respectively). Data represent three independent experiments (n=30). Error bars are SD, p values determined using ANOVA with Tukey’s post hoc test for significance; p <0.05 is significant.

### 5’ C-rich (ss)telomeric DNA at telomere-specific DSBs in G1 is not dependent on conventional exonucleases, nor does it engage in homology dependent repair

The presence of extensive tracks of ssDNA at telomeric DSBs in G1 suggested that long range resection was occurring. Therefore, we investigated the role of conventional end-processing exonucleases MRE11 (3’-to-5’) and EXO1 (5’-to-3’), known mediators of resection at genomic DSB sites and at telomeres, respectively, in phospho-RPA foci induction following ENT expression. Chemical inhibition of MRE11 (via treatment with the small molecule inhibitor PFM01) in EN-T expressing BJ1 hTERT G1 cells, did not significantly influence phospho-RPA32 foci, which were only slightly reduced compared to EN-T controls (p = 0.24 **Fig 5C**). Phospho-RPA32 foci were also only slightly reduced when BJ1 hTERT cells were partially depleted of EXO1 (siRNA); the difference was not statistically significant (p = 0.10 **Fig 5C**). Thus, the majority of observed resection at telomeric DSBs in G1 human cells appears to occur independently of conventional resection machinery.

An alternative explanation for the presence of ssDNA at telomeric DSBs in G1 could be that it represents an attempt to regenerate a normal (ss)telomeric G-rich overhang for T-loop formation and end-protection (18). The SNM1B/Apollo (5’-to-3’ exo) nuclease has been shown to be necessary for generation of the telomeric 3’ G-rich overhang at blunt-ended leading-strand telomeres in mice (54, 55). Therefore, we hypothesized that Apollo may act bidirectionally at telomeric DSBs, explaining the C-rich overhangs observed. However, EN-T-expressing Apollo^-/-^ EJ-30 human cells exhibited only a slight reduction in C-rich (ss)telomeric foci in G1 compared to wild type (WT) cells (p = 0.37, **Supp Fig 5A**). Additionally, EN-T expressing Apollo^-/-^ EJ-30 cells in G1 had more telomere phospho-RPA32 foci than EN-T expressing EJ-30 G1 WT cells (p = 0.099 **Supp Fig 5B**). Furthermore, both measures of telomeric ssDNA were slightly increased in the Apollo^-/-^ S/G2 populations compared to WT cells. Thus, the Apollo nuclease is not responsible for the extensive resection observed at telomeric DSBs in G1 human cells.

To determine whether telomere DSB-induced resected (ss)telomeric DNA in G1 could represent an early element of HR dependent repair (for strand-invasion), we also evaluated induction of RAD51 foci post EN-T transfection. While RAD51 foci were observed at telomeres in EN-T expressing EJ-30 S/G2 cells, they were *not* detected in EN-T expressing BJ1 hTERT or EJ-30 G1 cells (**Fig 6A**). Additionally, neither RAD52, nor repair associated DNA synthesis (BrdU incorporation) were detected following induction of telomeric DSBs in BJ1 hTERT G1 cells (**Fig 6B; Supp Fig 3C**). Taken together, these results indicate that telomeric ssDNA at telomeric DSBs in G1 human cells is not generated by conventional DSB or telomere resection machinery, nor does it in engage in resection-dependent recombinational repair, findings consistent with the majority of telomeric ssDNA in G1 being 5’ C-rich.

**Figure 6:**
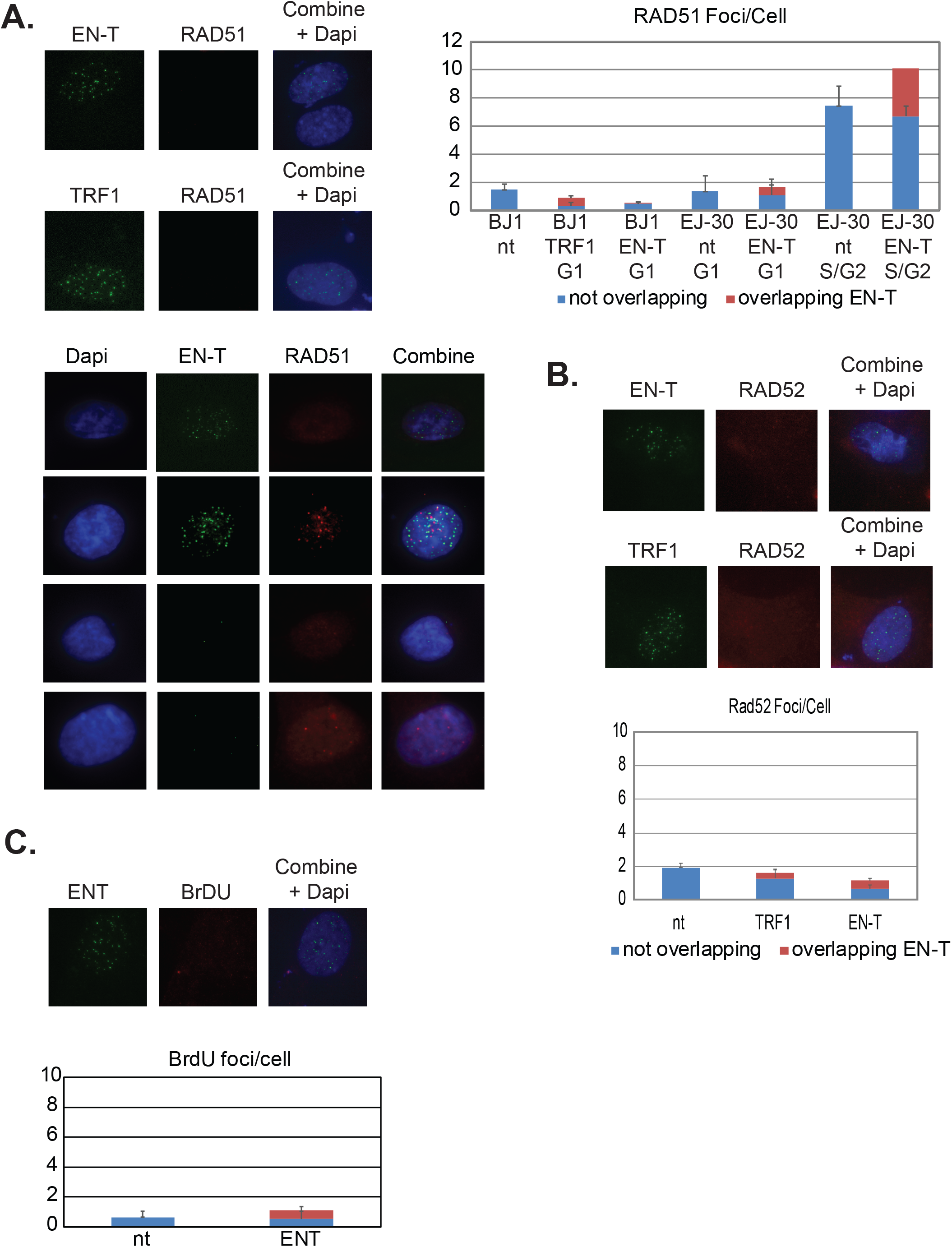
HR not occurring at telomeric DSBs in G1. **A.** Transfection of BJ1 hTERT or EJ-30 cells with EN-T did not induce RAD51 foci in G1. RAD51 foci were slightly increased in S/G2 EJ-30 cells following expression of EN-T, consistent with HR activity. **B**. Transfection of BJ1 hTERT cells with EN-T also did not induce RAD52 foci in G1. **C**. Repair associated DNA synthesis (BrdU incorporation) was also not detected in BJ1 hTERT G1 cells transfected with EN-T. Data represent three independent experiments (n= 30 BJ1 hTERT; n= 300 EJ-30 cells/experiment). Error bars are SD, p values determined using ANOVA with Tukey’s post hoc test for significance; p <0.05 is significant.

### 5’ C-rich (ss)telomeric DNA at telomere-specific DSBs in human ALT G1 cells is bound by telomeric RNA, TERRA

Telomeric C-rich (ss)overhangs [5’-CCCTAA-3’] are a previously proposed marker of the recombination dependent ALT pathway (40). Therefore, we evaluated whether telomeric DSBs in human U2OS (ALT) G1 cells (Supp Fig 2) resulted in significant increases in 5’ C-rich (ss)telomeric DNA. Considering the complementary nature of telomeric RNA, TERRA [5’-UUAGGG-3’], and that ALT cells possess higher levels of TERRA than non-ALT cells, we also monitored TERRA distribution specifically in U2OS G1 cells. A stable U2OS cell line expressing Geminin protein fused to Green Fluorescence Protein (Geminin-GFP) (49) was generated to positively identify, and eliminate from analyses, cells in G2; CENP-F staining also confirmed that Geminin-GFP positive cells were in G2. Cells negative for Geminin-GFP were in G1.

Utilizing native (non-denaturing) DNA FISH to detect 5’ C-rich (ss)telomeric DNA, no enrichment in EN-T and TRF1-only transfected FUCCI-U2OS G1 cells was observed (**Fig 7**). However, treatment with RnaseA and RnaseH (to remove TERRA) revealed a significant increase of resected (ss)telomeric C-rich DNA in cells transfected with EN-T, but not with TRF1-only. Together, these results suggest a protective role for telomeric RNA (TERRA) at resected 5’ C-rich (ss)telomeric DNA at telomeric DSB sites in G1 human ALT cells.

**Figure 7:**
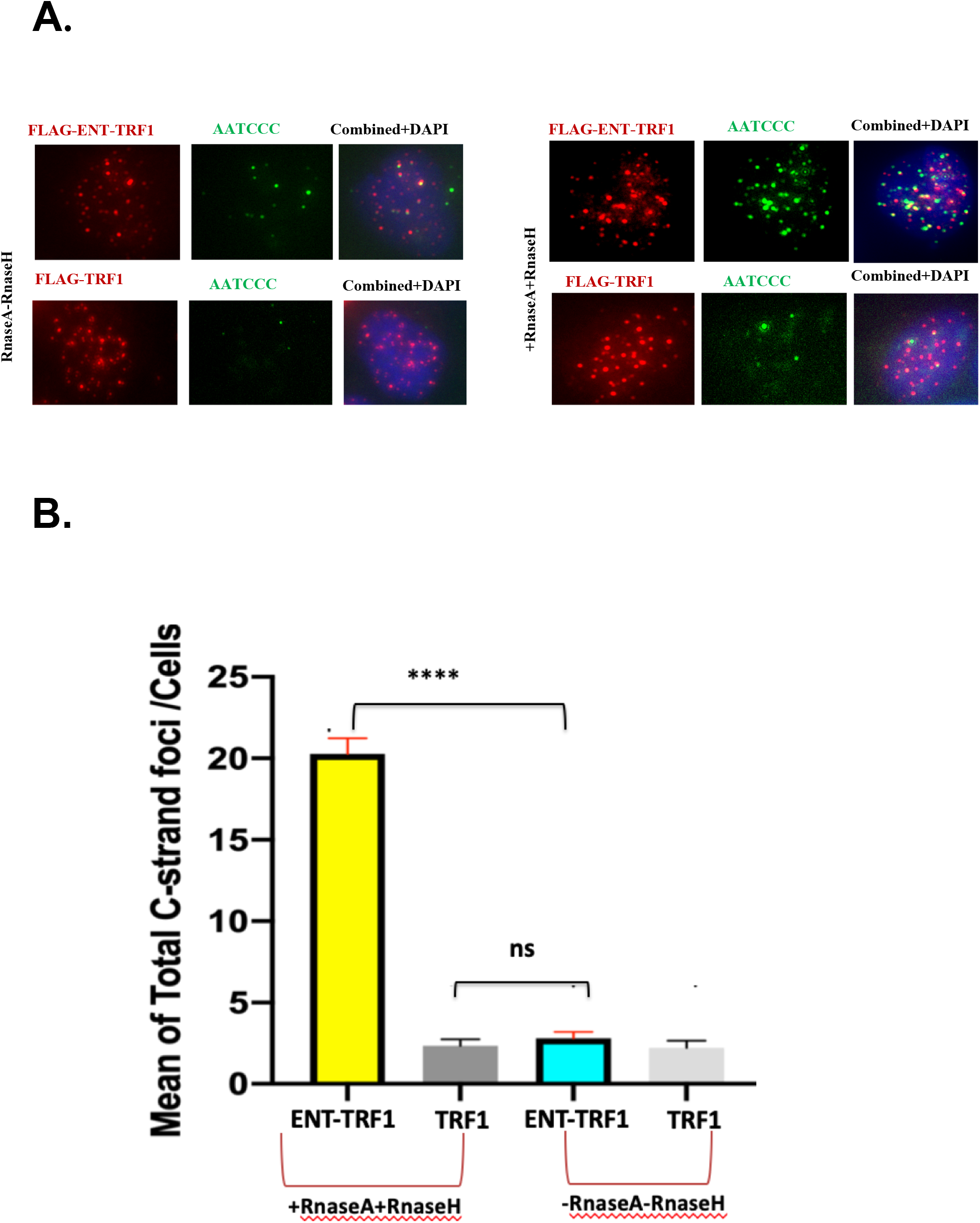
TERRA accumulates at telomeric 5’ C-rich (ss)overhangs in G1 U2OS cells. **A.)** Representative images of FUCCI-U2OS G1 cells transiently transfected with EN-T or TRF1-only (FLAG), labeled with G-rich telomere probe to detect C-rich (ss)telomeric DNA (AATCC), and merged views. Treatment with RNAseA and RNAseH removed telomeric RNA (TERRA) and revealed a significant increase in complementary 5’ C-rich (ss)telomeric DNA. **B.)** Quantification of average number telomeric C-rich foci. Data represents three experiments and values are expressed as SEM (*n* = 120-200). One-way ANOVA and Holm-Sidak Test significance; \*\*\**P* < 0.001; \*\**P* < 0.01; \**P* < 0.05; ns, not significant.

## DISCUSSION

Telomere-specific DSBs have generally been regarded as irreparable, as DDRs generated globally by ionizing radiation or other genotoxic agents fail to resolve when they occur at or near telomeres, and cells become senescent (31, 32). While repair of targeted telomeric DSBs has been observed in cycling cell populations, as well as specifically in S-phase, there is a dearth of evidence for DDRs or repair activity at telomeric DSBs in G1 cells (29, 30). To better understand human cellular responses to telomeric DSBs in G1, we investigated enzymatically induced (EN-T) telomere-targeted DSBs, specifically in telomerase positive BJ1hTERT (normal) and EJ-30 (cancer) cells, and in U2OS (ALT) cells.

Telomeric DSBs in G1 elicited early signatures of a DDR, as evidenced by γ-H2AX and MDC1 recruitment to telomere break sites. Notably however, while the commonly used DDR marker 53BP1 was recruited to telomeric DSBs in S/G2 – it was *not* present at those in G1 cells. Functionally, 53BP1 is most often associated with c-NHEJ, where it regulates 5’-to-3’ end-resection, but 53BP1 can also partially restrict resection during alt-NHEJ and HR repair (51, 56).

To gain mechanistic insight into this unexpected finding, we explored whether components of the telomere end-protection complex shelterin (19, 20) might be involved in thwarting 53BP1 recruitment to telomeric DSBs in G1 human cells. Telomere end-protection function was manipulated, without completely disrupting it, in an effort to alleviate inhibition of 53BP1 recruitment to telomeric DSBs while also avoiding dysfunctional telomere-induced foci (TIFs) (57). As near complete siRNA knockdown of TRF2 is necessary for a TIF response, we utilized an siRNA sequence that resulted in partial, sub-TIF-inducing depletion of TRF2, and combined it with EN-T or TRF1-only transfection in BJ1 hTERT cells. While partial TRF2 knockdown did not result in a TIF response in untransfected cells, it also did not alleviate inhibition of 53BP1 recruitment to telomeric DSBs in transfected G1 cells. Depletion of TRF2 in EN-T transfected EJ-30 cells also did not affect telomere fragmentation, indicating that TRF2 does not impact telomeric DSB repair. The absence of 53BP1 at telomeric DSBs in G1 was also observed in EN-T-transfected human cells depleted of another potential candidate, DNA-PKcs, previously shown to play a role in mammalian telomere end protection (58), and proposed to act in concert with TRF2 in preventing both c-NHEJ and alt-NHEJ at functional telomeres (59).

Compaction of telomeric chromatin has been proposed as a unifying physical mechanism by which shelterin protects telomeres from repair (21, 60). Therefore, we tested whether decompaction of genomic DNA could alleviate the repression of 53BP1 recruitment to telomeric DSBs in G1 human cells. Similar to partial TRF2 knockdown, treatment of EN-T transfected cells with a histone deacetylase inhibitor failed to result in recruitment of 53BP1 to G1 telomeric DSBs, suggesting that it may not be possible to relieve any potential influence of shelterin-mediated end-protection on inhibition of 53BP1 recruitment to telomeric DSBs in G1 without full deprotection of telomeres.

Consistent with the lack of 53BP1 at telomeric DSBs in G1 human cells, no evidence of c-NHEJ was observed, as neither shRNA depletion of DNA-PKcs nor chemical inhibition of DNA-PKcs catalytic activity influenced the response to EN-T-induced telomeric DSBs. Considering that both 53BP1 and c-NHEJ impede DSB repair associated DNA resection, we hypothesized that telomeric DSBs in G1 human cells may be especially vulnerable to resection. Indeed, pRPA coated (ss)telomeric DNA was detected following EN-T-mediated induction of telomeric DSBs in telomerase positive G1 cells, indicative of extensive resection at break sites; the detection limit of FISH is on the order of 0.5 Kb (61). Importantly and consistent with rapid truncation events and overall telomere shortening, telomeric DSBs in G1 human cells facilitated formation and enrichment of 5’ C-rich (ss)telomeric DNA, an observation supported by minimal dependence on MRE11, EXO1, or Apollo exonucleases. Given the abundance of (ss)telomeric DNA at broken telomeres, a potential role for resection-dependent repair was also interrogated, however no evidence of RAD51 (HR/BIR), RAD52 (BIR/SSA), or BrdU incorporation was observed. Alt-NHEJ was also not a likely candidate for telomeric DSB repair in G1, since it utilizes only a few base pairs of homology (~20) (62, 63), and is hindered by RPA binding to ssDNA (64).

Thus, although telomeric DSBs in G1 undergo extensive resection, they do not appear to be repaired in G1. One option may be that they attempt to reconstruct a 3’ G-rich (ss)overhang in order to form a protective T-loop and avoid c-NHEJ-mediated telomere-telomere fusion. Interestingly, both 3’ G-rich and 5’ C-rich telomeric overhangs have been proposed to mediate T-loop formation (40, 65). Therefore, resection may serve to stabilize broken telomeres during G1. This idea is supported by the fact that naturally shortened telomeres do not undergo fusion until nearly all telomeric repeats have been lost, suggesting that telomeres of nearly any length can be protected from repair activity (66). Further, (ss)telomeric overhangs at functional telomeres have been implicated in protection from repair (67). To extend this line of reasoning, resected telomeric DSBs may simply persist into S/G2, where telomeres critically shortened and/or rendered dysfunctional by internal DSBs could be elongated via telomerase-mediated or recombination-based ALT mechanisms, a notion consistent with telomeric 5’ C-rich (ss)overhangs as markers of the ALT pathway of telomere length maintenance (40, 68).

It also remains possible that some presently unappreciated pathway of repair operates at telomeric DSBs in human G1 cells. Potential candidates include RAD52-independent singlestrand annealing (SSA), as SSA was recently shown to take place in RAD52^-/-^ cells (69). Importantly however, RNA-templated DSB repair has recently been reported in human cells, a pathway that would be resection dependent and potentially not mediated by other conventional repair factors (70, 71). Our finding of telomeric RNA TERRA-bound 5’ C-rich (ss)telomeric DNA at telomeric DSB break sites in human ALT cells is particularly relevant in this regard. We propose that resected telomeric 5’ C-rich (ss)overhangs at telomeric DSBs in G1 human telomerase positive cells (with lower levels of TERRA) are coated primarily with RPA, presumably to further hamper NHEJ, while in G1 ALT cells (with higher levels of TERRA), transient telomeric RNA:DNA hybrids rapidly form to protect these exposed overhangs (41). Such dynamic interactions may well serve to preserve telomeric 5’ C-rich (ss)overhangs at telomeric DSB sites so that they persist into S/G2 phase (42) for replication (telomerase-mediated) or HR/template-dependent (ALT) elongation and restoration of functional telomeres, and in so doing they also present therapeutically relevant targets for disrupting telomere function and improving treatment of ALT positive tumors.

## MATERIALS AND METHODS

### Cell Culture and Transfections

Human U2OS (ALT), U2OS RAD52-YFP (obtained from Jiri Lucas, University of Copenhagen), and EJ-30 cancer (obtained from Dr. John Murnane, UCSF) cells were cultured in Dulbecco’s Modified Eagle Medium (DMEM, Hyclone) supplemented with 10% fetal bovine serum (FBS). Telomerase positive, apparently normal human BJ1 hTERT fibroblasts (ATCC), were cultured in Alpha-MEM (Hyclone) supplemented with 10% FBS.

TRAS1-EN-TRF1 and TRF1 plasmids, both driven by a CMV promoter and possessing a C-terminal Flag tag for visualization, were constructed from a CMV-driven TRAS1-EN-TRF1 plasmid obtained from Dr. Haruhiko Fujiwara (University of Tokyo). Transient transfections were carried out with Lipofectamine 3000 (Invitrogen) at 60-80% confluency in Opti-MEM (Gibco) for 20 minutes and replaced with normal media 8 hours later. Unless otherwise specified all experiments were carried out 48 hrs post transfection.

A U2OS cell line stably expressing FUCCI-Germinin (Green; S, G2/M) was established by transfecting cells with 0.5 ug of Kan-FUCCI-Green (S-G2-M) plasmid. Plasmids were delivered using Lipofectamine 3000 (Invitrogen) following the manufacturer’s instructions. Eight hours following transfection, Opti-MEM media was replaced with fresh DMEM media. One week later, cells were trypsinized and individual cells were seeded in a 96-well plate. A positive single clone was identified, and expanded in presence of 800 μg/ml G-418 sulfate (GoldBio). After reaching 90% confluency, cells were split into 24 well plate, 10 days later into 12 well plate, and a week after into 6 well plate. Cells were transferred into T-25 flask and then into a T-75 flask after 6 days. The DMEM media containing 800 μg/ml G-418 sulfate was changed every two days.

### Laser Micro-Irradiation

Laser micro-irradiations were performed with a Zeiss LSM880 confocal microscope using a 405nm laser at 100% with settings of 50 iterations and a 15 us pixel dwell. Spatially defined stripes of damage were generated through nuclei of cells followed by a recovery period of 30 min. Immunofluorescence and imaging of micro-irradiated cells were carried out as for other experiments and as described below.

### RNA interference

siRNA was initially delivered into cells using RNAiMAX in OptiMEM media according to manufacturer instructions, followed by replacement with normal media 5 hours later. 24 hours following initial siRNA delivery, cells were co-transfected with EN-T or TRF1-only and appropriate siRNA in Lipofectamine 3000 according to manufacturer instructions, and then fixed or harvested 48hrs later. siRNA sequences were as follows: TRF2: 5’-GAGGAUGAACUGUUUCAAGdtdt-3’ (anti-sense also included 3’ dtdt), EXO1: 5’-UGCCUUUGCUAAUCCAAUCCCACGC-3’.

### Inhibitors

BJ1hTERT fibroblasts were treated with either a kinase inhibitor that prevents DNA-PKcs autophosphorylation (NU7026, Sigma) or MRE11 endonuclease activity inhibitor (PFM01, Thermofisher). NU7026 was used at a concentration of 10uM for 24 hours prior to harvesting cells as per previous (72). Alternatively, PFM01 was used at a concentration of 100uM for 8 hours preceding fixation. For chromatin relaxation, cells were treated with trichostatin A (TSA, Sigma) at the specified concentrations for 24 hrs prior to cell fixation.

### Western Blotting

Cell pellets were washed in phosphate buffered saline (PBS), and then incubated in lysis buffer for 10 minutes. Lysis buffer consisted of Mammalian Protein Extraction Reagent (M-PER, ThermoFisher) with protease inhibitors (complete mini EDTA free, Sigma-Aldrich), and in cases when phosphorylated proteins were being detected, phosphatase inhibitors (PhosSTOP, Sigma-Alrdrich). Following isolation of protein, the Bradford assay was used to quantify protein (BioRad). 30ug of protein was loaded into precast SDS-PAGE gels (Mini-Protean TGX, 4-15%, BioRad) in Tris/Glycine/SDS buffer followed by electrophoretic separation for roughly 1.5 hours at 125V. After electrophoresis, proteins were transferred to a polyvinylidene fluoride (PVDF) membrane in Tris/Glycine buffer with 10-15% methanol for 16-20 hr at 30V at 4°C. An even protein transfer was verified by reversibly staining membranes with Poncaeu S solution (Sigma-Aldrich, 0.1% w/v in 1% acetic acid). Next, membranes were blocked in 5% non-fat dry milk (NFDM), or bovine serum albumin (BSA) in 1X Tris buffered saline with 0.1% Tween 20 (TBST) from 30 min to 1 hr with gentle shaking. Blocking solution was then replaced with fresh blocking solution containing the appropriate dilution of primary antibody and incubated from 2 hr to overnight with gentle shaking. Following primary antibody incubation, membranes were washed in 1X TBST for 4 washes of 10 min each with gentle shaking. Next, fresh blocking solution was added with the appropriate dilution of a horseradish peroxidase (HRP) labeled secondary antibody and incubated from 2 to 4 hr followed by another series of 4 washes in 1X TBST. Following the final wash, membranes were rinsed in PBS. To visualize proteins, membranes were treated with SuperSignal™ West Pico Chemiluminescent Substrate according to the manufacturer instructions (ThermoFisher) and imaged on a ChemiDoc™ XRS+ imager with ImageLab^™^ software (BioRad).

Primary antibodies for western blotting included Rabbit Anti-phospho serine2056 DNA-PKcs (Abcam ab1249181, 1:2000), Mouse Anti-DNA-PKcs (ThermoFisher MS-423-P, 1:10000), Mouse Anti-TRF2 (SantaCruz sc-271710, 1:500), Mouse Anti-phospho serine1981 ATM (Upstate 05-740, 1:1000), Rabbit Anti-phospho Thr68 CHK2 (Cell signaling 2661, 1:1000), Rabbit Anti-EXO1 (Proteintech 16352-1-AP, 1:500). HRP labeled secondary antibodies included Donkey Anti-Rabbit (Jackson ImmunoResearch 711-035-152, 1:20000), and Rabbit Anti-Mouse (ThermoFisher 816720, 1:10000).

### Immunofluorescence

Unless stated otherwise, cells were grown on chamber slides, rinsed twice in PBS, fixed in freshly prepared 4% paraformaldehyde (PFA) for 10 min at room temperature, and then permeabilized in 0.2% Triton X-100 in PBS for 4-10 min. Next, cells were blocked in 10% normal goat serum (NGS), or 5% BSA in 1xPBS for 40 min and then incubated with primary antibodies diluted in blocking solution for 1 hr at 37°C or overnight at 4°C. Following primary incubations cells were washed 3 times in 1xPBS at 42°C. After washes cells were incubated with fluorophore-conjugated goat secondary antibodies for 30 min at 37°C. Finally, cells were washed again as before and counterstained with Prolong Gold Antifade reagent with DAPI (Invitrogen).

Primary antibodies and concentrations included: Rabbit Anti-53BP1 (Bethyl A300-272A, 1:800), Rabbit Anti-γ-H2AX (Bethyl A300-081, 1:1000), Mouse Anti-Flag (Sigma M2 F1804, 1:2000-4000), Rabbit Anti-RPA70 (Cell signaling #2267, 1:50), Rabbit Anti-phospho S4/S8 RPA32 (Bethyl A300-245A 1:2000), Mouse Anti-gammaH2AX (Millipore 05-636, 1:1500), Rabbit Anti-Cyclin A (Santa Cruz SC-751, 1:500), Rabbit Anti-MDC1 (Bethyl A300-051A, 1:1000), Rabbit Anti-RAD51 (H-92 SC-8349, 1:800), Sheep Anti-RAD52 (kind gift from Jiri Lukas Lab, 1:100), Rat anti-BrdU (BioRad OBT0030, 1:200), Rabbit Anti-phospho S15 p53 (Abcam Ab18128-50, 1:500)

Secondary antibodies and concentrations included: Alexa-488 Goat anti-Mouse (ThermoFisher A11029, 1:750), Alexa-594 Goat anti-Mouse (ThermoFisher A11005, 1:750), Alexa-647 Goat anti-Mouse (ThermoFisher A21235, 1:350), Alexa-488 Donkey anti-Mouse (ThermoFisher 21202, 1:750), Alexa-488 Goat anti-Rabbit (ThermoFisher A11008, 1:750), Alexa-594 Goat anti-Rabbit (ThermoFisher A11012, 1:750), Alexa-555 Goat anti-Rat (ThermoFisher A21434, 1:750), Alexa-647 Donkey anti-Sheep (ThermoFisher A21448, 1:350), Alexa-488 Donkey anti-Mouse (ThermoFisher A21202 1:750).

### BrdU Incorporation Assay

Cells were pulse-labeled with the thymidine analog Bromodeoxyuridine / 5-bromo-2’-deoxyuridine (BrdU; ThermoFisher) for 2 hr (50mM) and then fixed for 15 min in 4% PFA at room temperature. Next, cells were permeabilized for 20 min with 0.1% Triton x-100 in PBS, followed by DNA denaturation for 10 min on ice with 1N HCl and then 10 min at room temperature with 2N HCL. Cells were then washed with phosphate citric acid buffer pH 7.4 for 10 min at room temperature. Finally, cells were washed for 5 min in permeabilization solution. Blocking was then carried out for 30 min at 37°C in 5% NGS with 0.1% Triton X-100 in PBS. Antibody incubations, washing steps, and counterstaining were carried out as described for immunofluorescence.

### Non-denaturing Immuno-FISH

Combined immunofluorescence and fluorescence in-situ hybridization (FISH) experiments were carried out on cells grown on chamber slides. Cells were initially fixed in 4% PFA for 5 min at room temperature. Next, cells were permeabilized for 4 min in 0.2% Triton X-100 in PBS. Following permeabilization, cells were blocked and immunostained as described in the immunofluorescence section. After the last washing step, cells were post-fixed in 4% PFA for 15 min at room temperature. Next, cells were dehydrated in an ethanol series (75%, 85%, 95%) for 2 min each and allowed to air dry. While slides were air drying, the hybridization solution was prepared by combining 36ul formamide, 12ul 0.05M Tris-HCL, 2.5ul 0.1M KCL, 0.6 ul 0.1M MgCl2 and 0.5ul 0.5uM Peptide Nucleic Acid (PNA) telomere probe (TelC-Alexa488 or TelG-Cy3, Biosynthesis) in 20% acetic acid. Hybridization solution was then denatured at 85°C for 10 min followed by cooling on ice. After cooling, 50ul of hybridization solution was added to each slide, then slides were coverslipped, and incubated at 37°C in a humidified chamber for 6 hr. Following hybridization, coverslips were removed and slides washed twice in 50% formamide in 2X SSC (2.5 min @ 42°C), twice in 2X SSC (2.5 min @ 42°C) and twice in 2X SSC + 0.1% NP-40 (2.5 minutes 42°C). Following the final wash, cells were counterstained with Prolong Gold Antifade with DAPI.

### Fluorescence Microscopy and Image Analysis

Images were acquired using a Zeiss Axio Imager.Z2 epi-fluorescent microscope using a 63X/1.4 N.A. objective (Plan-APOCHROMAT, Zeiss). For the majority of targets, images were blindly and subjectively thresholded and segmented followed by determination of foci overlap (50% overlap scored as positive) in Metamorph 7.7 (Molecular Devices). For RAD52-YFP, RPA and phospho-RPA foci analysis, cells tended to have very few or an abundance of foci and scoring was therefore done on the basis of whether a cell had >4 foci overlapping Flag.

Analysis of the laser microirradiation experiment involved first thresholding TRF2 foci using a fixed value for all images. Next, these thresholded foci were converted to regions in Metamorph and these regions transferred to γ-H2AX or 53BP1 images. Next, the average intensity within the transferred regions was compared to that within pseudo-random regions of comparable dimensions generated by rotating TRF2 images by 90°.

For BrdU foci analysis in BJ1 hTERTs, untransfected S-phase cells were excluded from analysis, which were identified by very bright pan nuclear staining.

For DAPI-intensity based cell cycle analysis, DAPI intensity was collected alongside foci by subjective thresholding and segmentation in Metamorph followed by histogram generation. Foci counts were sorted based on whether the nuclear intensity fell into the clear G1 peak or the S/G2 tail. The border region between cell cycle phases of four DAPI intensity bins was excluded from analysis to ensure accurate classification of cells.

### Telomere Restriction Fragment (TRF) Southern Blots

The TRF assay was performed using the TeloTAGGG^™^ southern blotting kit (Roche) according to the manufacturer instructions with some modifications, including a longer probe hybridization time (6hr), as well as a longer incubation time with Anti-DIG antibody (1hr). 2ug of sample DNA were loaded per lane and blots were imaged on a ChemiDoc^™^ XRS+ imager with ImageLab™ software (BioRad). Quantitation of mean TRF length was performed using TeloTool software according to the manufacturer’s protocol.

### Statistical analyses

EN-T validation experiments in U20S cells were done in duplicate (50 cells per replicate). Experiments with BJ1 hTERT cells involved three independent experiments for each condition with at least 30 cells per replicate for imaging experiments. The exception to this was micro-irradiation experiments, which were done in duplicate with 15 cells per replicate. Experiments with EJ-30 cells were also done in triplicate; however, the number of cells imaged totaled at least 300 per condition to allow for DAPI intensity histogram generation.

Error bars on bar graphs represent standard deviations, and p-values were computed when experiments were done in triplicate, and are provided in the text when less than 0.05 (significance threshold). When two groups were being compared, p-values were generated via students T-tests, alternatively, when three or more groups were being compared an ANOVA with a Tukey’s post hoc test was used. ANOVAs were either one way or two way depending on the number of categorical independent variables.

## CONFLICT OF INTEREST

The authors declare that there are no conflicts of interest.

## ACKNOWLEDGEMENTS

The authors sincerely thank Dr. Haruhiko Fujiwara for generously supplying the TRAS1-EN-TRF1 and TRF1 plasmids, and Dr. Jiri Lucas for U2OS cell lines. We also gratefully acknowledge funding from NASA (NNX14AB02G).

---

**Supplemental Figure 1:**
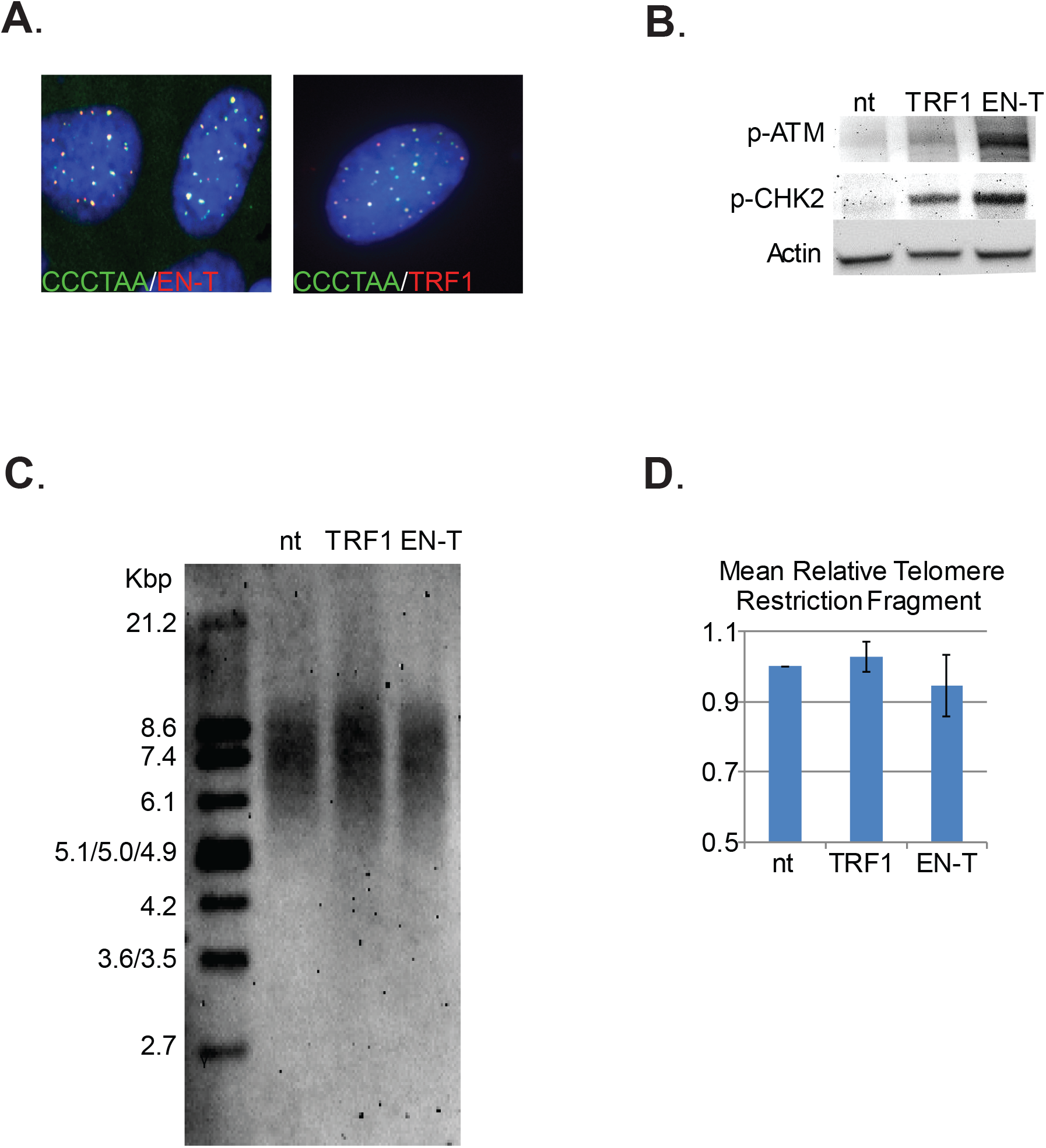
Characterization of telomere-specific cutting endonuclease TRAS-ENT (ENT). **A.** Overexpressed EN-T or TRF1-only co-localized with telomere repeats in U20S, EJ-30, and BJ1 hTERT cells (shown). **B.** Expression of EN-T in EJ-30 cells activated DDR signaling, evidenced by P-S1981-ATM and P-Thr68-CHK2. **C.** Expression of EN-T in EJ-30 cells also resulted in fragmentation of telomeric DNA on southern blot of telomeric restriction fragments (TRF); **D.** quantification of non-transfected (nt), TRF1 control, and ENT transfected cells. Data represent three independent experiments, with n=50 (U2OS), n=30 (BJ1 hTERT) or n=300 (EJ-30) cells/experiment. Error bars are SD, p-values < 0.05, significant.

**Supplemental Figure 2:**
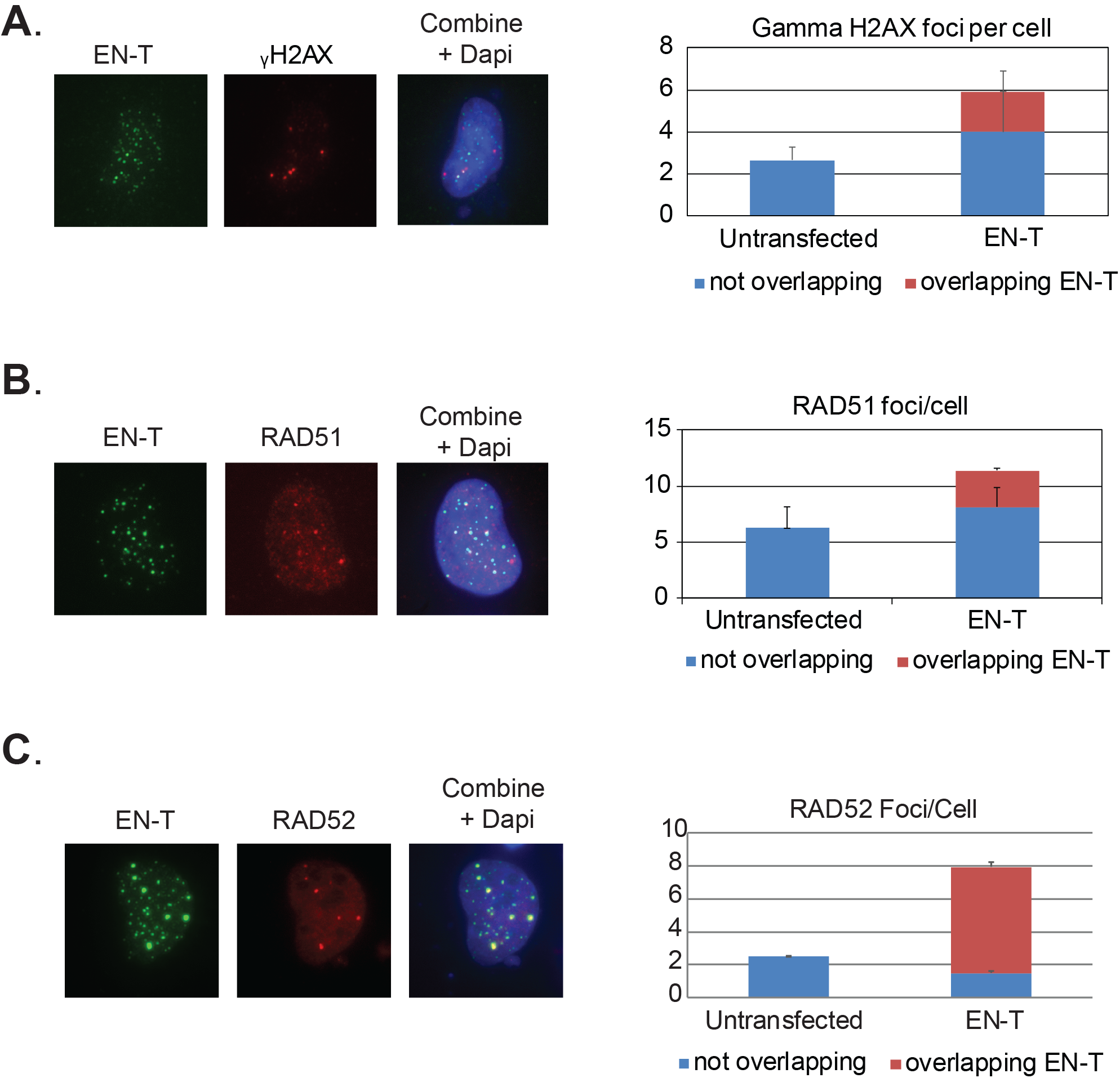
Additional characterization of EN-T system. **A.** Transfection of cycling U20S (ALT) cells with EN-T triggered a telomeric DDR in terms of γ-H2AX foci, which frequently overlapped with ENT-flag (red bar). **B, C.** Induced telomeric DSBs in cycling U20S cells also stimulated recruitment of RAD51 and RAD52, mediators of HR and BIR respectively, both of which frequently overlapped with ENT-flag.

**Supplemental Figure 3:**
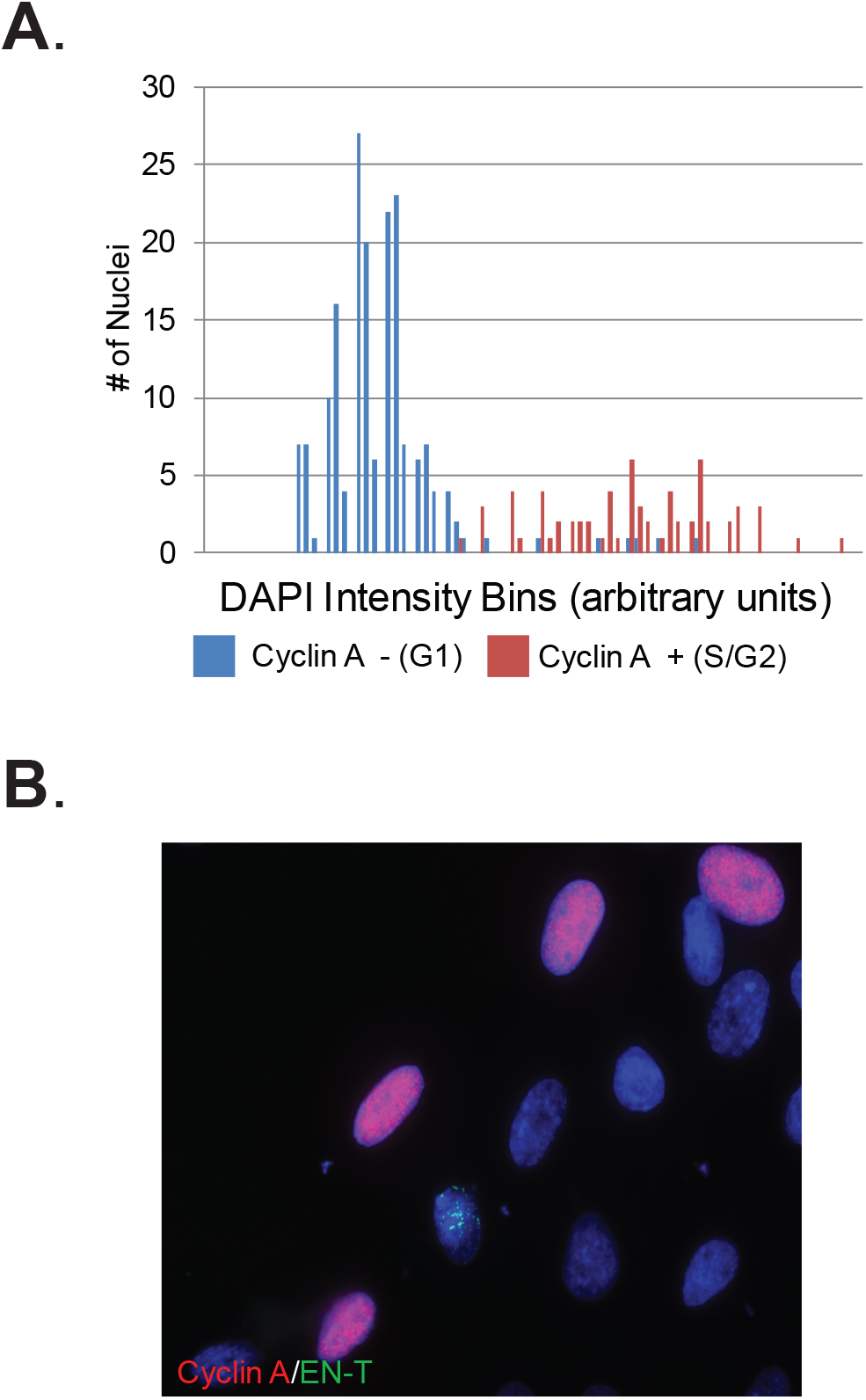
DAPI intensity histograms to identify cells in G1. **A.** DAPI intensity histograms were generated from 63x images of approximately 300 cells per experiment. Exclusion of Cyclin A from the low DAPI intensity peak region of the histogram (blue bars) verified that these cells were in G1 phase of the cell cycle; data shown represent merged histograms from 3 replicates totaling 300 EJ-30 cells. **B.** DAPI intensity histograms were not necessary for identification of BJ1 hTERT G1 cells, as EN-T and TRF1-only transfected cells were almost exclusively negative for Cyclin A, consistent with the vast majority of transfected BJ1 hTERT cells being in G1 phase 48 hr post transfection when analyses were done. Image illustrates that while the population of cells contains many cyclin A positive cells (red), the relatively few transfected cells (green foci; EN-T) were always cyclin A negative (in G1). Additionally, BrdU incorporation was not detected in BJ hTERT cells transfected with EN-T (Figure 6).

**Supplemental Figure 4:**
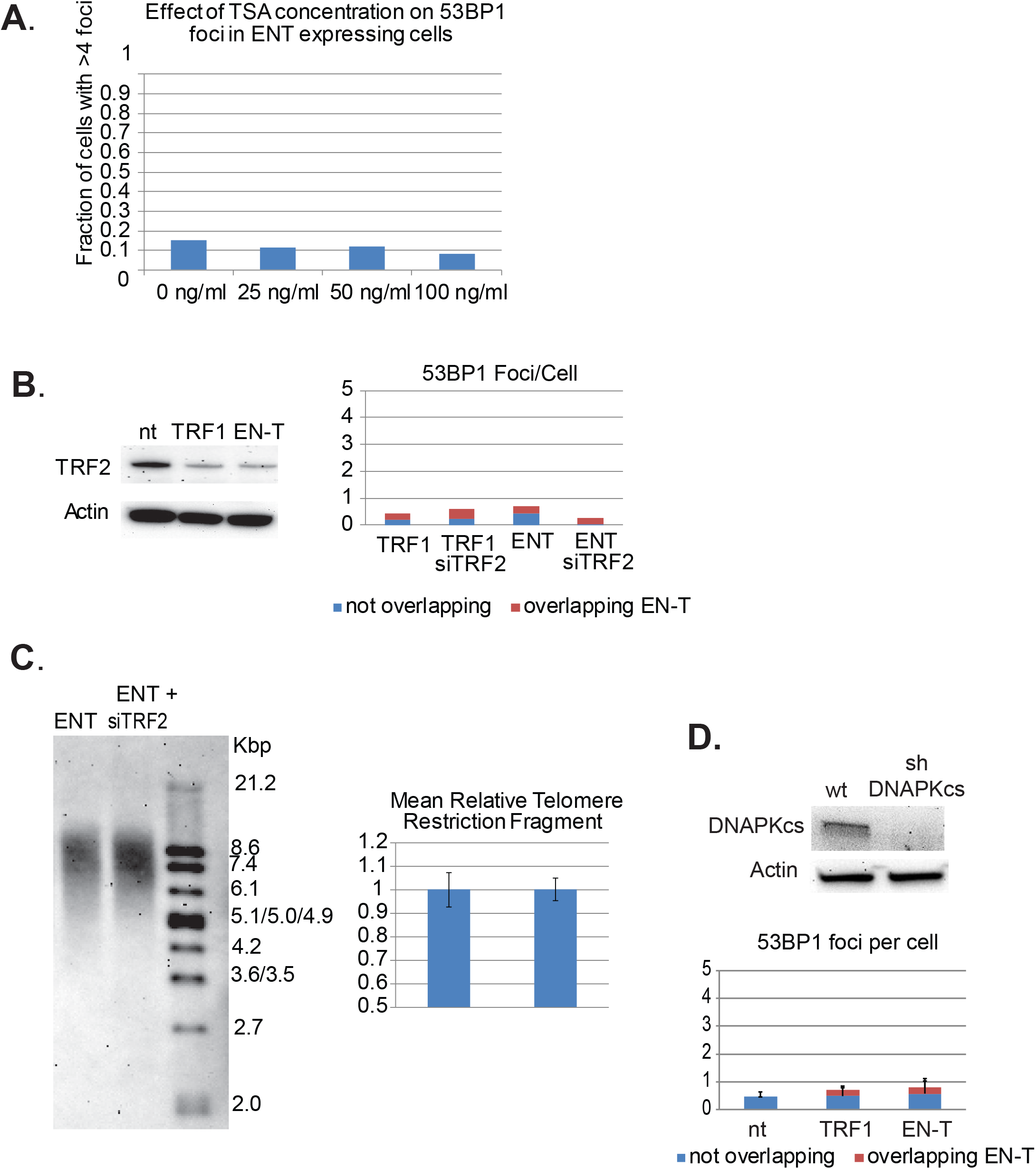
Compromised telomeric end-protection does not promote 53BP1 recruitment to broken telomeres. **A.** Relaxation of chromatin via treatment with trichostatin A (TSA) did not result in 53BP1 foci induction in EN-T expressing cells at any concentration. **B.** Partial depletion (siRNA knockdown) of TRF2 did not influence induction of 53BP1 foci in EN-T or TRF1-only transfected BJ1hTERT cells. **C.** siRNA knockdown of TRF2 also had no measurable effect on the degree of telomere fragmentation in EJ-30 cells transfected with EN-T. **D.** shRNA knockdown of DNA-PKcs did not promote 53BP1 recruitment to telomeric DSBs in EN-T transfected BJ1-hTERT cells.

**Supplemental Figure 5:**
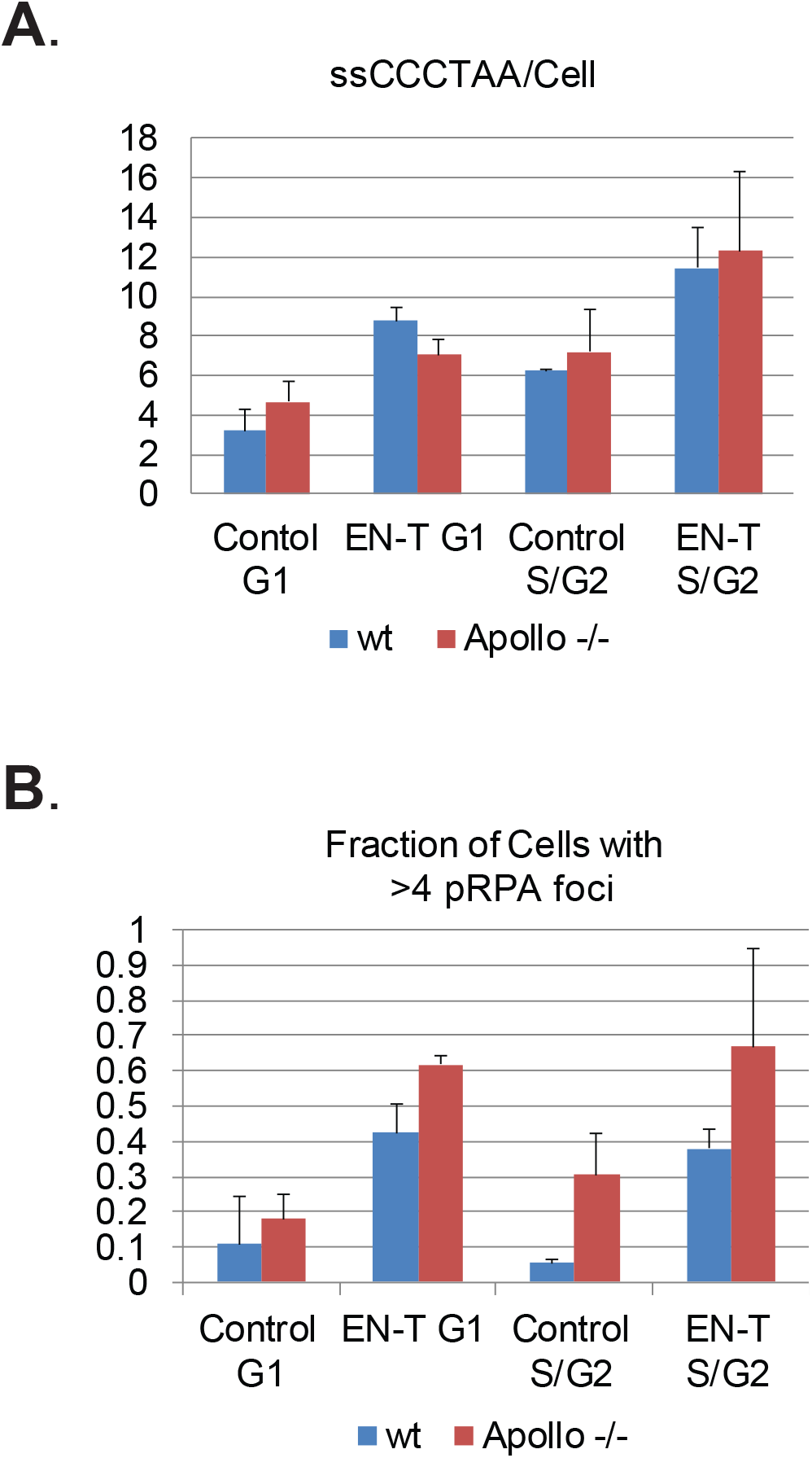
Role of Apollo endonuclease in the generation of ssDNA at telomeric DSBs. **A.** While telomeric ssDNA (5’-CCCTAA-3’) was slightly reduced in EN-T expressing EJ-30 Apollo^-/-^ G1 cells relative to EN-T expressing control (wild type) EJ-30 cells, **B.** phospho-RPA32 foci were somewhat increased. Additionally, both telomeric ssDNA and phospho-RPA32 foci were increased in EN-T expressing EJ-30 Apollo^-/-^ S/G2 cells.

## REFERENCES

1. Meyne J, Ratliff RL, Moyzis RK. Conservation of the human telomere sequence (TTAGGG)n among vertebrates. Proc Natl Acad Sci USA. 1989;86(18):7049–53.

2. Azzalin CM, Reichenbach P, Khoriauli L, Giulotto E, Lingner J. Telomeric repeat containing RNA and RNA surveillance factors at mammalian chromosome ends. Science (New York, NY). 2007;318(5851):798–801.

3. Arora R, Lee Y, Wischnewski H, Brun CM, Schwarz T, Azzalin CM. RNaseH1 regulates TERRA-telomeric DNA hybrids and telomere maintenance in ALT tumour cells. Nature communications. 2014;5:5220.

4. Balk B, Maicher A, Dees M, Klermund J, Luke-Glaser S, Bender K, et al. Telomeric RNA-DNA hybrids affect telomere-length dynamics and senescence. Nature structural & molecular biology. 2013;20(10):1199–205.

5. Bettin N, Oss Pegorar C, Cusanelli E. The Emerging Roles of TERRA in Telomere Maintenance and Genome Stability. Cells. 2019;8(3).

6. Deng Z, Norseen J, Wiedmer A, Riethman H, Lieberman PM. TERRA RNA binding to TRF2 facilitates heterochromatin formation and ORC recruitment at telomeres. Molecular cell. 2009;35(4):403–13.

7. Montero JJ, Lopez-Silanes I, Megias D, M FF, Castells-Garcia A, Blasco MA. TERRA recruitment of polycomb to telomeres is essential for histone trymethylation marks at telomeric heterochromatin. Nature communications. 2018;9(1):1548.

8. Makarov VL, Hirose Y, Langmore JP. Long G tails at both ends of human chromosomes suggest a C strand degradation mechanism for telomere shortening. Cell. 1997;88(5):657–66.

9. Greider CW, Blackburn EH. Identification of a specific telomere terminal transferase activity in Tetrahymena extracts. Cell. 1985;43(2 Pt 1):405–13.

10. Kim NW, Piatyszek MA, Prowse KR, Harley CB, West MD, Ho PL, et al. Specific association of human telomerase activity with immortal cells and cancer. Science. 1994;266(5193):2011–5.

11. Batista LFZ. Telomere biology in stem cells and reprogramming. Prog Mol Biol Transl Sci. 2014;125:67–88.

12. Bryan TM, Englezou A, Dalla-Pozza L, Dunham MA, Reddel RR. Evidence for an alternative mechanism for maintaining telomere length in human tumors and tumor-derived cell lines. Nat Med. 1997;3(11):1271–4.

13. Henson JD, Cao Y, Huschtscha LI, Chang AC, Au AYM, Pickett HA, et al. DNA C-circles are specific and quantifiable markers of alternative-lengthening-of-telomeres activity. Nat Biotechnol. 2009;27(12):1181–5.

14. Cesare AJ, Reddel RR. Alternative lengthening of telomeres: models, mechanisms and implications. Nat Rev Genet. 2010;11(5):319–30.

15. Bailey SM, Brenneman MA, Goodwin EH. Frequent recombination in telomeric DNA may extend the proliferative life of telomerase-negative cells. Nucleic Acids Res. 2004;32(12):3743–51.

16. Murnane JP, Sabatier L, Marder BA, Morgan WF. Telomere dynamics in an immortal human cell line. EMBO J. 1994;13(20):4953–62.

17. Heaphy CM, Subhawong AP, Hong SM, Goggins MG, Montgomery EA, Gabrielson E, et al. Prevalence of the alternative lengthening of telomeres telomere maintenance mechanism in human cancer subtypes. The American journal of pathology. 2011;179(4):1608–15.

18. Griffith JD, Comeau L, Rosenfield S, Stansel RM, Bianchi A, Moss H, et al. Mammalian telomeres end in a large duplex loop. Cell. 1999;97(4):503–14.

19. de Lange T. Shelterin: the protein complex that shapes and safeguards human telomeres. Genes Dev. 2005;19(18):2100–10.

20. de Lange T. How Shelterin Solves the Telomere End-Protection Problem. Cold Spring Harbor Symposia on Quantitative Biology. 2010;75(0):167–77.

21. Baker AM, Fu Q, Hayward W, Victoria S, Pedroso IM, Lindsay SM, et al. The telomere binding protein TRF2 induces chromatin compaction. PloS One. 2011;6(4):e19124.

22. Sfeir A, de Lange T. Removal of shelterin reveals the telomere end-protection problem. Science (New York, NY). 2012;336(6081):593–7.

23. de Lange T. A loopy view of telomere evolution. Frontiers in Genetics. 2015;6:321.

24. van Steensel B, Smogorzewska A, de Lange T. TRF2 Protects Human Telomeres from End-to-End Fusions. Cell. 1998;92(3):401–13.

25. Bae NS, Baumann P. A RAP1/TRF2 complex inhibits nonhomologous end-joining at human telomeric DNA ends. Molecular cell. 2007;26(3):323–34.

26. Kibe T, Osawa GA, Keegan CE, de Lange T. Telomere protection by TPP1 is mediated by POT1a and POT1b. Molecular and Cellular Biology. 2010;30(4):1059–66.

27. Anzai T, Takahashi H, Fujiwara H. Sequence-specific recognition and cleavage of telomeric repeat (TTAGG)(n) by endonuclease of non-long terminal repeat retrotransposon TRAS1. Molecular and Cellular Biology. 2001;21(1):100–8.

28. Yoshitake K, Aoyagi H, Fujiwara H. Creation of a novel telomere-cutting endonuclease based on the EN domain of telomere-specific non-long terminal repeat retrotransposon, TRAS1. Mobile DNA. 2010;1(1):13.

29. Mao P, Liu J, Zhang Z, Zhang H, Liu H, Gao S, et al. Homologous recombination-dependent repair of telomeric DSBs in proliferating human cells. Nature communications. 2016;7:12154.

30. Doksani Y, de Lange T. Telomere-Internal Double-Strand Breaks Are Repaired by Homologous Recombination and PARP1/Lig3-Dependent End-Joining. Cell Reports. 2016;17(6):1646–56.

31. Hewitt G, Jurk D, Marques FDM, Correia-Melo C, Hardy T, Gackowska A, et al. Telomeres are favoured targets of a persistent DNA damage response in ageing and stress-induced senescence. Nature communications. 2012;3:708.

32. Fumagalli M, Rossiello F, Clerici M, Barozzi S, Cittaro D, Kaplunov JM, et al. Telomeric DNA damage is irreparable and causes persistent DNA-damage-response activation. Nature Cell Biology. 2012;14(4):355–65.

33. Lieber MR. The Mechanism of Double-Strand DNA Break Repair by the Nonhomologous DNA End-Joining Pathway. Annual Review of Biochemistry. 2010;79(1):181–211.

34. Chapman JR, Taylor MRG, Boulton SJ. Playing the end game: DNA double-strand break repair pathway choice. Molecular cell. 2012;47(4):497–510.

35. Chang HHY, Pannunzio NR, Adachi N, Lieber MR. Non-homologous DNA end joining and alternative pathways to double-strand break repair. Nature Reviews Molecular Cell Biology. 2017;18(8):495–506.

36. Miller D, Reynolds GE, Mejia R, Stark JM, Murnane JP. Subtelomeric regions in mammalian cells are deficient in DNA double-strand break repair. DNA repair. 2011;10(5):536–44.

37. Muraki K, Han L, Miller D, Murnane JP. The role of ATM in the deficiency in nonhomologous end-joining near telomeres in a human cancer cell line. PLoS genetics. 2013;9(3):e1003386.

38. Maréchal A, Zou L. RPA-coated single-stranded DNA as a platform for post-translational modifications in the DNA damage response. Cell Research. 2015;25(1):9–23.

39. Lam YC, Akhter S, Gu P, Ye J, Poulet A, Giraud-Panis M-J, et al. SNMIB/Apollo protects leading-strand telomeres against NHEJ-mediated repair. EMBO J. 2010;29(13):2230–41.

40. Oganesian L, Karlseder J. Mammalian 5’ C-rich telomeric overhangs are a mark of recombination-dependent telomere maintenance. Mol Cell. 2011;42(2):224–36.

41. Ohle C, Tesorero R, Schermann G, Dobrev N, Sinning I, Fischer T. Transient RNA-DNA Hybrids Are Required for Efficient Double-Strand Break Repair. Cell. 2016;167(4):1001–13.e7.

42. D’Alessandro G, Whelan DR, Howard SM, Vitelli V, Renaudin X, Adamowicz M, et al. BRCA2 controls DNA:RNA hybrid level at DSBs by mediating RNase H2 recruitment. Nature communications. 2018;9(1):5376-.

43. Zschenker O, Kulkarni A, Miller D, Reynolds GE, Granger-Locatelli M, Pottier G, et al. Increased sensitivity of subtelomeric regions to DNA double-strand breaks in a human cancer cell line. DNA repair. 2009;8(8):886–900.

44. Munoz P, Blanco R, de Carcer G, Schoeftner S, Benetti R, Flores JM, et al. TRF1 controls telomere length and mitotic fidelity in epithelial homeostasis. Molecular and cellular biology. 2009;29(6):1608–25.

45. Palm W, de Lange T. How shelterin protects mammalian telomeres. Annual review of genetics. 2008;42:301–34.

46. van Steensel B, de Lange T. Control of telomere length by the human telomeric protein TRF1. Nature. 1997;385(6618):740–3.

47. Cho NW, Dilley RL, Lampson MA, Greenberg RA. Interchromosomal homology searches drive directional ALT telomere movement and synapsis. Cell. 2014;159(1):108–21.

48. Dilley RL, Verma P, Cho NW, Winters HD, Wondisford AR, Greenberg RA. Break-induced telomere synthesis underlies alternative telomere maintenance. Nature. 2016;539(7627):54–8.

49. Sakaue-Sawano A, Kurokawa H, Morimura T, Hanyu A, Hama H, Osawa H, et al. Visualizing spatiotemporal dynamics of multicellular cell-cycle progression. Cell. 2008;132(3):487–98.

50. Stewart GS, Wang B, Bignell CR, Taylor AMR, Elledge SJ. MDC1 is a mediator of the mammalian DNA damage checkpoint. Nature. 2003;421(6926):961–6.

51. Xiong X, Du Z, Wang Y, Feng Z, Fan P, Yan C, et al. 53BP1 promotes microhomology-mediated end-joining in G1-phase cells. Nucleic Acids Research. 2015;43(3):1659–70.

52. Dimitrova N, Chen Y-CM, Spector DL, Lange T. 53BP1 promotes non-homologous end joining of telomeres by increasing chromatin mobility. Nature. 2008;456(7221):524–8.

53. Zimmermann M, Lottersberger F, Buonomo SB, Sfeir A, de Lange T. 53BP1 regulates DSB repair using Rif1 to control 5’ end resection. Science. 2013;339(6120):700–4.

54. Wu P, van Overbeek M, Rooney S, de Lange T. Apollo contributes to G overhang maintenance and protects leading-end telomeres. Molecular cell. 2010;39(4):606–17.

55. Wu P, Takai H, de Lange T. Telomeric 3’ overhangs derive from resection by Exo1 and Apollo and fill-in by POT1b-associated CST. Cell. 2012;150(1):39–52.

56. Ochs F, Somyajit K, Altmeyer M, Rask M-B, Lukas J, Lukas C. 53BP1 fosters fidelity of homology-directed DNA repair. Nature structural & molecular biology. 2016;23(8):714–21.

57. Cesare AJ, Hayashi MT, Crabbe L, Karlseder J. The telomere deprotection response is functionally distinct from the genomic DNA damage response. Molecular cell. 2013;51(2):141–55.

58. Bailey SM, Meyne J, Chen DJ, Kurimasa A, Li GC, Lehnert BE, et al. DNA double-strand break repair proteins are required to cap the ends of mammalian chromosomes. Proc Natl Acad Sci USA. 1999;96(26):14899–904.

59. Bombarde O, Boby C, Gomez D, Frit P, Giraud-Panis M-J, Gilson E, et al. TRF2/RAP1 and DNA-PK mediate a double protection against joining at telomeric ends. EMBO J. 2010;29(9):1573–84.

60. Bandaria JN, Qin P, Berk V, Chu S, Yildiz A. Shelterin Protects Chromosome Ends by Compacting Telomeric Chromatin. Cell. 2016;164(4):735–46.

61. Ramakrishnan S, Sulochana KN. Manual of Medical Laboratory Techniques: Jaypee Brothers Medical Publishers Pvt. Ltd.; 2012 2012/12/15/. 453 p.

62. Truong LN, Li Y, Shi LZ, Hwang PY-H, He J, Wang H, et al. Microhomology-mediated End Joining and Homologous Recombination share the initial end resection step to repair DNA double-strand breaks in mammalian cells. Proceedings of the National Academy of Sciences of the United States of America. 2013;110(19):7720–5.

63. Symington LS, Gautier J. Double-Strand Break End Resection and Repair Pathway Choice. Annual Review of Genetics. 2011;45(1):247–71.

64. Deng SK, Gibb B, de Almeida MJ, Greene EC, Symington LS. RPA antagonizes microhomology-mediated repair of DNA double-strand breaks. Nature structural & molecular biology. 2014;21(4):405–12.

65. Verdun RE, Karlseder J. The DNA damage machinery and homologous recombination pathway act consecutively to protect human telomeres. Cell. 2006;127(4):709–20.

66. Capper R, Britt-Compton B, Tankimanova M, Rowson J, Letsolo B, Man S, et al. The nature of telomere fusion and a definition of the critical telomere length in human cells. Genes & Development. 2007;21(19):2495–508.

67. Gong Y, de Lange T. A Shld1-controlled POT1a provides support for repression of ATR signaling at telomeres through RPA exclusion. Molecular cell. 2010;40(3):377–87.

68. Oganesian L, Karlseder J. 5’ C-rich telomeric overhangs are an outcome of rapid telomere truncation events. DNA Repair (Amst). 2013;12(3):238–45.

69. Kan Y, Batada NN, Hendrickson EA. Human somatic cells deficient for RAD52 are impaired for viral integration and compromised for most aspects of homology-directed repair. DNA repair. 2017;55:64–75.

70. Meers C, Keskin H, Storici F. DNA repair by RNA: Templated, or not templated, that is the question. DNA repair. 2016;44:17–21.

71. Mazina OM, Keskin H, Hanamshet K, Storici F, Mazin AV. Rad52 Inverse Strand Exchange Drives RNA-Templated DNA Double-Strand Break Repair. Molecular cell. 2017;67(1):19–29.e3.

72. Le PN, Maranon DG, Altina NH, Battaglia CLR, Bailey SM. TERRA, hnRNP A1, and DNA-PKcs Interactions at Human Telomeres. Frontiers in Oncology. 2013;3:91.

